# Decoupling Quantification and Identification for Label-Free Top-Down Proteomics Improves Proteome Coverage

**DOI:** 10.64898/2026.11.06.764767

**Authors:** Philipp T. Kaulich, Christian Treitz, Tom D. Müller, Josefine Rost, Oliver Kohlbacher, Kyowon Jeong, Andreas Tholey

## Abstract

Quantifying proteoforms is essential for understanding biological processes. The most widely used quantification approach in top-down proteomics (TDP) is label-free quantification (LFQ), where quantification and identification compete for acquisition time. We identified missing values, insufficient fragmentation, and undersampling of low-abundance proteoforms as major factors limiting conventional TDP-LFQ workflows. Our proposed workflow decouples quantification and identification through MS1-only quantification followed by targeted data-dependent MS2 identification. The MS1-only acquisition more than doubled the number of mass features quantified reliably and improved quantification precision. Focusing targeted MS2 acquisition on significantly differentially abundant features enables optimization of MS2 parameters and increases identification coverage by 50%. The workflow was validated across different chromatographic conditions, MS acquisition setups, and sample preparation strategies. In *Escherichia coli* grown on glucose or acetate, the increased depth enabled by this workflow revealed biological insights inaccessible to a previously published bottom-up proteomics dataset. Overall, decoupling quantification and identification improves quantitative depth and proteoform coverage in TDP-LFQ and provides a flexible strategy for high-sensitivity and high-throughput proteoform quantification.

**TOC:** 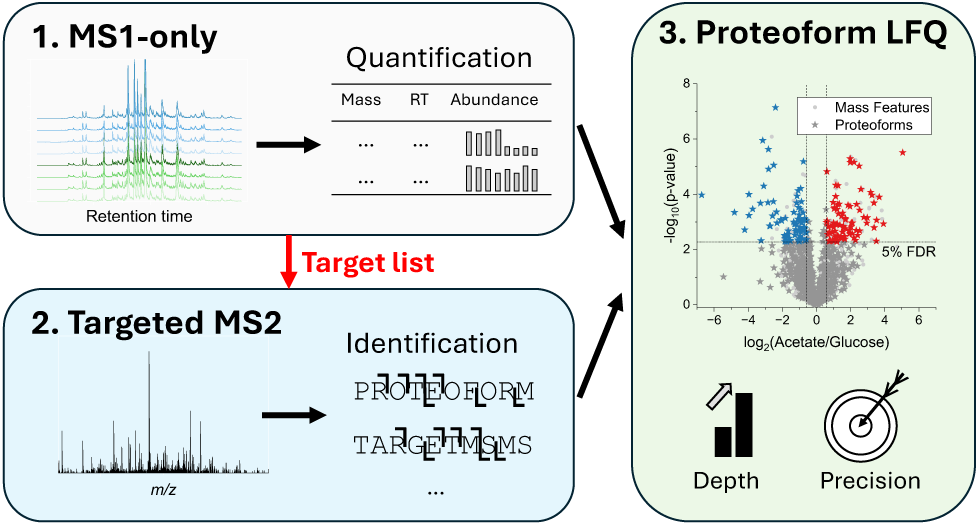

## 1 Introduction

Proteoforms, defined as the distinct molecular forms in which proteins are present, represent the functional units of biological systems.^1^ Since different proteoforms of a given protein, arising from genetic variations, alternative splicing, or post-translational modifications,^2^ can fulfill distinct biological functions, their characterization is essential for understanding cellular molecular mechanisms. In contrast to bottom-up proteomics (BUP), top-down proteomics (TDP) analyzes intact proteoforms from complex samples and, thus, enables the elucidation of proteome complexity. While proteoform identification determines which molecular species are present in a sample, quantitative analysis provides another layer of information, such as how proteoform abundances change across biological conditions.

Several strategies have been developed for proteoform quantification, including label-based and label-free approaches. Label-based quantification strategies, including non-isobaric and isobaric approaches, are readily compatible with multidimensional separation schemes and enable multiplexing.^3–5^ However, they typically require expensive reagents, have limited multiplexing capacity, and may introduce artificial modifications from incomplete or excessive labeling. In contrast, in label-free quantification (LFQ) workflows, all samples are analyzed individually by LC-MS, followed by spectral deconvolution and mass feature quantification.^6^ A mass feature is defined by its deconvolved monoisotopic mass, elution profile, and abundance, with the abundance corresponding to the integrated intensity across the elution profile of all observed charge states and isotopes.^7^ The mass features are then assigned to proteoforms, and relative proteoform abundance between conditions is determined by statistical analysis of their abundances across samples. Since LFQ is experimentally straightforward, cost-effective, and easily scalable to larger sample numbers, it is currently the most widely used quantification strategy in TDP.^8–11^ Furthermore, recent advances in acquisition and data analysis have substantially improved mass feature detection and quantification, e.g., through MS1-only acquisition^12^ and approaches for resolving overlapping proteoform signals.^13,14^

However, current TDP-LFQ workflows remain limited in analytical depth and quantitative precision, since MS1-based quantification of intact proteoforms is challenged by many factors stemming from spectral complexity, including overlapping charge-state envelopes, co-eluting species, and the low reproducibility in detection of low-abundance mass features across LC-MS runs. High analytical sensitivity is essential for comprehensive proteoform quantification because many biologically relevant proteoforms occur at low abundance. Classical strategies to increase analysis depth, such as multidimensional separation, require additional sample preparation steps that may introduce technical variation and reduce quantitative reproducibility.^6^ Thus, approaches are required that improve quantitative performance without requiring additional sample preparation steps.

To address the limitations in quantification and breadth of coverage in TDP, we developed a dual-step LFQ workflow to improve quantitative depth and precision. Using *Escherichia coli* grown on glucose or acetate as a model system, we first identified missing values, insufficient fragmentation, and intensity-dependent undersampling of low-abundance proteoforms as the major factors limiting conventional data-dependent acquisition (DDA)-based TDP-LFQ workflows. Based on these findings, we decoupled mass feature quantification from proteoform identification: First, MS1-only acquisitions were performed for mass feature quantification and statistical analysis, followed by targeted data-dependent MS2 acquisition for proteoform identification. This design enables independent optimization of MS1 and MS2 parameters, resulting in a doubling of the number of quantified mass features and the identification of approximately 50% more significantly differentially abundant proteoforms. The workflow was validated across different MS setups, chromatographic gradient lengths, and sample preparation strategies. Furthermore, we compared TDP-based proteoform-level quantification with a reanalyzed BUP dataset, revealing biological insights masked by protein-level data.

## 2 Experimental Section

Based on the limitations of conventional TDP-LFQ, we developed a dual-step workflow that separates MS1-only quantification from targeted MS2 proteoform identification. First, MS1-only measurements are performed across all samples to quantify mass features and identify significantly differentially abundant features. Subsequently, targeted MS2 measurements are performed to identify the proteoforms corresponding to these selected features. Separating quantification and identification allows independent optimization of MS1 and MS2 acquisition parameters. An overview of the workflow is provided in **Figure 1**.

**Figure 1:**
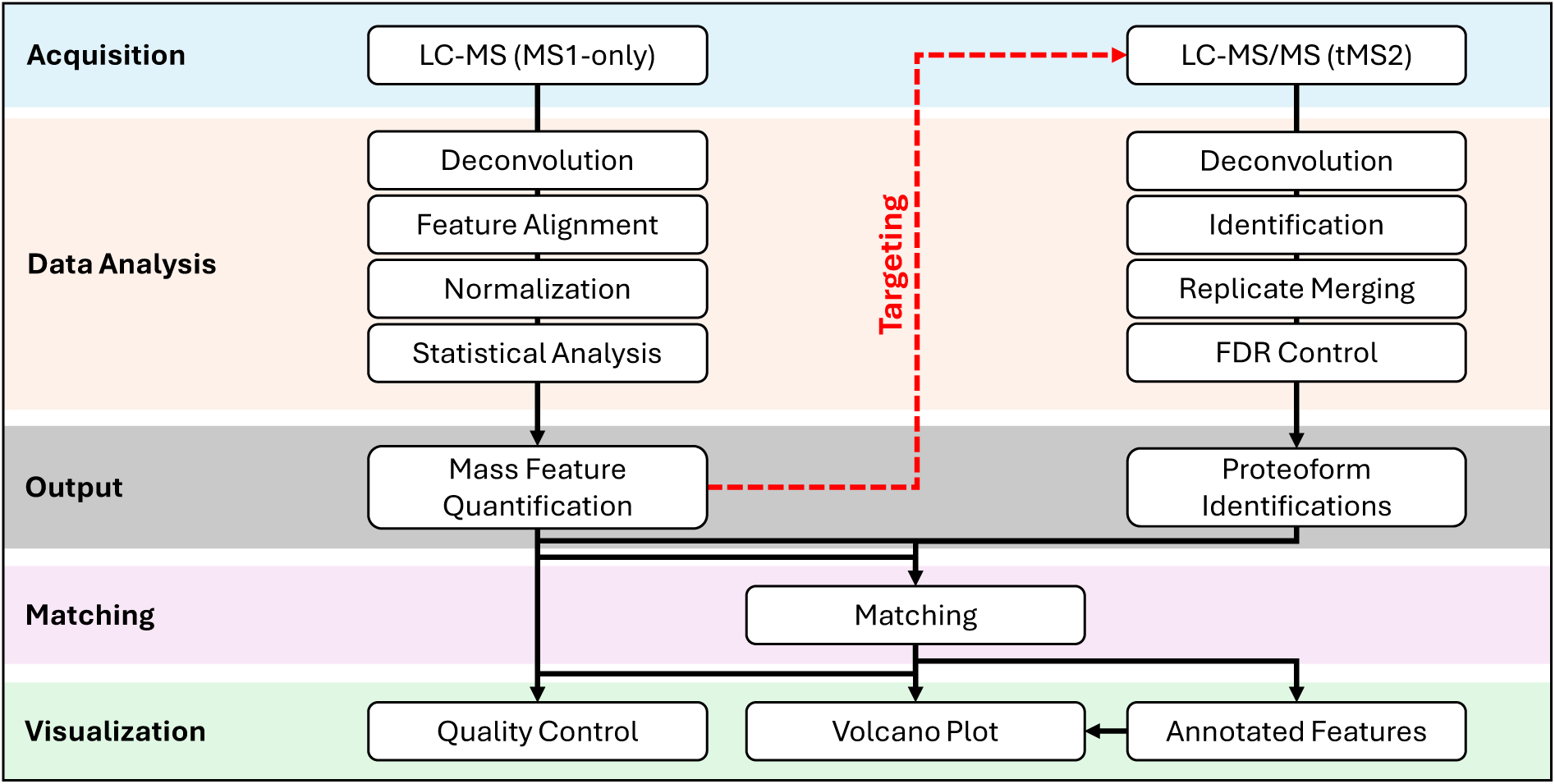
Overview of the dual-step MS1-only/targeted MS2 quantification strategy for TDP-LFQ. First, mass feature detection is performed by MS1-only acquisition, followed by deconvolution, feature alignment, normalization, and statistical analysis. Quantified mass features are subsequently used to generate inclusion lists for targeted data-dependent MS2 (tMS2) acquisition based on retention time, *m/z,* and charge state. After tMS2 measurements, proteoforms are identified by deconvolution and database search. Identifications from various files are combined (replicate merging and 1% target-decoy FDR control), and the identified proteoforms are matched to quantified mass features based on accurate mass and retention time tags.

### 2.1 Materials

All chemicals and solutions were purchased from Sigma-Aldrich (Steinheim, Germany) unless stated otherwise. The cOmplete protease inhibitor was from Roche (Basel, Switzerland). Ultrapure Milli-Q water (18.2 MΩ⋅cm) was prepared using an Arium VF 611 system (Sartorius, Göttingen, Germany).

### 2.2 *Escherichia coli* Cultivation and Cell Lysis

*Escherichia coli* K-12 strain MG1655 was cultivated as previously described.^15^ In brief, biological replicates of bacteria were cultured in M9 minimal medium^16^ supplemented either with glucose (15 mM) or acetate (45 mM) as the sole carbon source (four replicates per condition). After inoculating the cells at an optical density (OD600) of 0.1, they were incubated at 37 °C under constant shaking, and OD600 was measured regularly. Once the cell culture reached an OD600 of 1, the cells were harvested by centrifugation for 3 min at 3,000 rcf and at room temperature. The cells were washed with ultrapure water and stored at −80 °C prior to cell lysis.

For cell lysis and protein extraction, the cell pellets were resuspended in 1 ml 6.4 M guanidinium hydrochloride, 200 mM triethylammonium bicarbonate (TEAB, pH 8.5), 1× cOmplete protease inhibitor. Lysis was performed by freeze-thaw cycling (5 cycles: 3 min at −70 °C in an ethanol bath, 2 min in an ultrasonication bath at room temperature). After centrifugation at 21,100 rcf for 25 min (4 °C), the supernatant was collected, and the protein concentration was determined using the Pierce BCA Protein Assay Kit (Thermo Fisher). The extracted proteins were stored at −80 °C until further use.

### 2.3 Reduction & Alkylation

For proteoform reduction and alkylation, 1 mg of proteoforms per replicate was diluted to 330 µl with ultrapure water. Reduction was performed with tris(2-carboxyethyl)phosphine at a final concentration of 2.5 mM (40 min, 37 °C, 800 rpm), and alkylation with iodoacetamide at a final concentration of 5 mM (30 min, 20 °C, 800 rpm, in the dark). For proteoform purification, 100 µl (∼300 µg) was aliquoted and subjected to methanol-chloroform-water precipitation.^17^ In brief, 400 µl of methanol and 100 µl of chloroform were added to the sample, followed by vortexing. Subsequently, 300 µl of water was added, the sample was vortexed again, and incubated for 2 min at room temperature. Phase separation was induced by centrifugation (5 min, 14,000 rcf, room temperature), and the upper phase was carefully removed. Then, 400 µl of methanol was added to the sample and mixed by inversion. After centrifugation (5 min, 21,100 rcf, room temperature), the supernatant was discarded, and the resulting protein pellet was washed with methanol.

For resolubilization,^18^ the pellet was dissolved in 30 µl of pre-cooled (−20 °C) 80% (v/v) formic acid, briefly vortexed, and incubated for 5 min at −20 °C. Subsequently, 270 µl of water was added (final protein concentration ∼1 mg/ml), and the samples were stored at −80 °C until LC-MS/MS analysis.

### 2.4 Solid-phase Extraction

Solid-phase extraction (SPE) using a C18 SepPak (1 cc, 50 mg; Waters, Milford, MA, USA) was performed to enrich suitable proteoforms for TDP (smaller than approximately 30 kDa), as previously described,^19^ with minor modifications. Briefly, 1 mg of *Escherichia coli* proteoforms per replicate was adjusted to a final volume of 800 µl using 100 mM TEAB (pH 8.5). Prior to sample loading, the SepPak cartridge was conditioned with 2× 1 ml acetonitrile and equilibrated with 2× 1 ml 100 mM TEAB. The cartridge was washed with 2× 1 ml 100 mM TEAB before proteoforms were eluted using 150 µl each of 70% acetonitrile in 30 mM TEAB and 100% acetonitrile. The eluate was lyophilized, and the proteoforms were resuspended in 200 µl of 3% acetonitrile, 0.1% trifluoroacetic acid, and stored at −80 °C prior to LC-MS analysis.

### 2.5 LC-MS Analysis

Prior to mass spectrometric analysis, proteoforms were separated using a Vanquish Neo UHPLC system (Thermo Fisher Scientific) equipped with either a C4 (15 cm x 75 µm, 2.5 µm, 150 Å, Thermo Fisher Scientific) or an MAbPac column (15 cm × 150 µm, 4 µm, 1,500 Å; Thermo Fisher Scientific). For sample loading, a C4 precolumn (C4 PepMap300, 5 µm, 300 Å, Thermo Fisher Scientific) was utilized. For 30 min gradients, 0.5 µl per sample was injected; for 60 min gradients, 1 µl, and for 90 min gradients, 3 µl. Proteoform separation was performed using eluent A (0.05% formic acid) and eluent B (80% acetonitrile, 0.04% formic acid). The flow rate was set to 300 nl/min for the C4 column and 1.5 µl/min for the MAbPac column at 45 °C. Proteoforms were separated using a 30-min, 60-min, or 90-min linear gradient from 25% to 60% eluent B.

The LC was coupled online to an Orbitrap Eclipse Tribrid mass spectrometer (Thermo Fisher Scientific) operated with the Thermo Scientific Orbitrap Tribrid Series instrument control application, v4.2.4321. For measurements performed with high-field asymmetric waveform ion mobility spectrometry (FAIMS), the FAIMS Pro Duo Interface was installed and operated at standard resolution (100 °C), 4.6 L/min total carrier gas flow, and a single compensation voltage of −40 V. Acquisition was performed using an ion-routing multipole pressure of 0.008 Torr (“peptide mode”).

MS1 acquisition parameters were optimized based on the number, reproducibility, and quantitative quality of detected mass features (**Supplementary Figures 1 and 2**): MS1 spectra were acquired from 500-1,850 *m/z* at a resolution of 120,000 with an automatic gain control (AGC) target of 1,000%, a maximum injection time of 246 ms, and 1 microscan.

**Figure 2:**
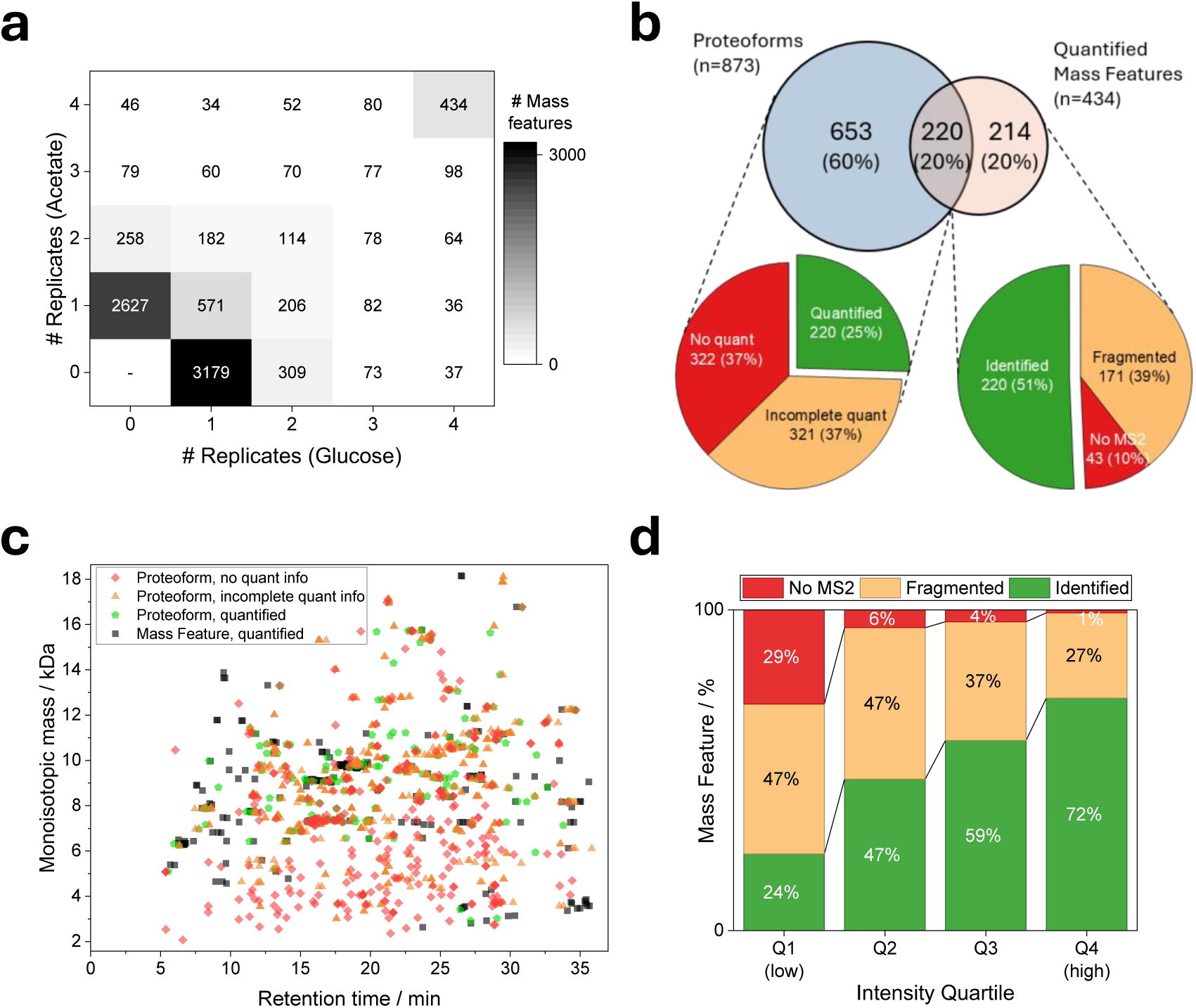
Major limitations of the conventional DDA-based MS1/MS2 TDP-LFQ approach. **a** Distribution of mass features according to the number of biological replicates in each condition in which they were detected. **b** Overlap between identified proteoforms and quantified mass features (Venn diagram), with major reasons why identified proteoforms could not be quantified (left pie chart), or why quantified mass features could not be assigned to proteoform identifications (right pie chart). **c** Scatter plot of quantified mass features and identified proteoforms by precursor mass and retention time. **d** Effect of mass feature intensity on precursor selection for MS2 fragmentation and subsequent proteoform identification. Quantified: proteoforms assigned to mass features quantified in all replicates. Incomplete quantification: proteoforms assigned to mass features detected in at least one, but not all, replicates. No quantification: proteoforms not assigned to any quantified mass feature. Identified: quantified mass features assigned to a proteoform identification. Fragmented: quantified mass features selected for MS2 acquisition but not yielding a confident proteoform identification. No MS2: quantified mass features not selected for MS2 fragmentation.

For classical data-dependent MS1/MS2 experiments, the MS2 spectra (200-2,000 *m/z*, 60,000 resolution, 1,000% AGC target, 118 ms maximum injection time, 2 microscans) were acquired within a 1.5 s (30 min gradient) or 3 s (60 min gradient) cycle time, and the most intense precursor ions (charge states 4-50, dynamic exclusion enabled with n=2 within 30 s, 60 s exclusion, ±1 *m/z* tolerance) were fragmented using an isolation window of 2 *m/z* and high-energy collision-induced dissociation (HCD) with a normalized collision energy (NCE) of 30%.

### 2.6 Targeted Data-dependent Acquisition

For targeted data-dependent MS2 acquisition of quantified or significantly differentially abundant mass features, inclusion lists were generated specifying precursor average mass, charge state range, retention time, retention time window, and intensity threshold. Charge state ranges were determined from the deconvoluted MS1-only data by selecting the most abundant charge state together with the ±1 and ±2 adjacent charge states (based on FLASHDeconv output). The target retention time corresponded to the apex retention time of the respective deconvolved mass features. Furthermore, a retention time window of 2 min and an intensity threshold of 1×10e4 were applied for all targets. MS1 parameters were the same as described above. If no precursor (i.e., charge state *z* within *m/z*±0.5) from the inclusion list was detected in an MS1 scan, the most intense ions were selected for fragmentation (dynamic exclusion enabled). The optimized MS2 parameters for the targeted acquisition were: 120,000 resolution, 246 ms maximum injection time, and two fragment scans per precursor, with HCD fragmentation (NCE 30%) and collision-induced dissociation (CID, NCE 35%). For targeted MS2 analyses, a pooled sample combining all four biological replicates from both conditions was analyzed. To identify condition-specific proteoforms, one randomly selected biological replicate per condition was analyzed instead.

### 2.7 Data Analysis

#### 2.7.1 Proteoform Quantification and Identification

For mass feature deconvolution and quantification, several deconvolution algorithms were evaluated, including FLASHDeconv (OpenMS-3.1.0-pre-FVdeploy-2025-09-25),^14^ FLASHQuant (OpenMS-3.1.0-pre-FVdeploy-2025-09-19),^13^ TopFD (TopPIC Suite v1.7.10),^7^ and Xtract (within ProSightPD v4.2) (**Supplementary Figure 3**).^20^ Based on the overall number of detected mass features (detected in at least one replicate), quantified mass features (detected in all replicates), the number of significantly differentially abundant mass features (q <5%, absolute log_2_ fold change >0.58), and quantitative reproducibility, FLASHDeconv showed the best overall performance and was therefore selected for all subsequent analyses. Default parameters were used, except that the minimum precursor mass and minimum charge state were set to 2,000 Da and +4. FLASHDeconv was used to generate both deconvolved mass feature tables and corresponding msAlign files for downstream proteoform identification.

Proteoform identification was performed using TopPIC (v1.7.10).^21^ Protein sequences from the *Escherichia coli* strain K12 were downloaded from UniProt^22^ (taxon ID: 83333; release 2025_02) and used as the reference FASTA database. The msAlign files generated by FLASHDeconv were used as input for database searching, and the following search parameters were applied: no fixed modifications; variable modifications included oxidation (+15.995 Da) on methionine residues and phosphorylation (+79.966 Da) on serine, threonine, and tyrosine residues; a maximum of three variable modifications per proteoform; and a maximum of one unexpected modification. The allowed precursor mass shift range was set to ±500 Da. Searches were performed in target-decoy mode, and all decoys were retained for subsequent FDR filtering.

Proteoform identifications from multiple LC-MS runs were combined as previously reported.^23^ In brief, all TopPIC identification lists were merged into a single list (including both target and decoy identifications). Proteoforms without unexpected modifications were combined if they had identical protein sequences, variable PTMs, and terminal modifications. For proteoforms containing unexpected modifications, identifications with the same protein accession, proteoform sequence, and variable PTMs were grouped and additionally required to match within a mass and retention time tolerance of 30 ppm and 120 s. For each resulting group, the identification with the lowest (i.e., the best) E-value was retained. Then, all identifications were sorted by E-value in ascending order. Starting with the best-scoring identification, the cumulative numbers of target and decoy identifications were determined and used to estimate the false discovery rate (FDR) using the target-decoy approach. Only target proteoforms passing this global 1% FDR threshold were retained for downstream analysis.

An accurate mass and retention time tag strategy was applied for proteoform quantification:^24,25^ proteoforms were matched to quantified mass features with a precursor mass tolerance of 30 ppm and a retention time (in the case of mass features, apex retention time) tolerance of 120 s. To avoid ambiguous assignments, a strict one-to-one mapping strategy was applied. If multiple proteoform identifications matched the same quantified mass feature, only the identification with the smallest precursor mass deviation was retained. Likewise, if a proteoform identification matched multiple quantified mass features, it was assigned exclusively to the mass feature with the smallest precursor mass deviation. A proteoform was considered significantly differentially abundant if it was assigned to a significantly differentially abundant mass feature.

#### 2.7.2 Statistical Analysis

Deconvolved mass features from all LC-MS runs were aligned using a greedy clustering algorithm with a precursor mass tolerance of 20 ppm and a retention time tolerance of 120 s (feature alignment). Mass features were sorted by mass and were iteratively assigned to existing clusters if their precursor masses and retention times matched the current cluster centroid (mean precursor mass and mean retention time) within 20 ppm and 120 s; otherwise, a new cluster was initialized. After each assignment, the cluster centroid was recalculated using all mass features assigned to the cluster. The final list contained one cluster per aligned mass feature together with its corresponding intensities across all LC-MS runs.

Prior to statistical analysis, feature intensities were median-normalized. For each run, all intensities were multiplied by the ratio of the global median intensity across all runs to the median intensity of the respective run. After this, mass features were filtered to exclude those with missing values; i.e., only mass features with valid quantitative values in all replicates across both experimental conditions were retained for downstream statistical analysis.

Differential abundance between conditions was assessed using two-tailed Welch’s t-test on log_2_-transformed feature intensities. Resulting p-values were adjusted for multiple hypothesis testing using the Benjamini-Hochberg procedure. Mass features with an adjusted p-value (q-value) <0.05 and an absolute log₂ fold change >0.58 (corresponding to a >1.5-fold difference) were considered significantly differentially abundant.

#### 2.7.3 Biological Interpretation and Comparison with Bottom-Up Proteomics Data

To assess possible condition-dependent proteoforms in the biological interpretation of the data, a second statistical evaluation was performed, including mass features with valid quantitative values in all four replicates in one of the two cultivation conditions. Data imputation was performed, replacing missing values from a normal distribution (width 0.4, down-shift 2), and the same statistical testing described above was performed. Whenever non-imputed data were available, reported fold changes are solely based on these data.

For comparison with bottom-up proteomics, a publicly available 2D-LC MS isobaric-labeling-based quantitative dataset was reanalyzed (ProteomeXchange: PXD003863).^15,26^ The raw data were searched against the same *E. coli* database using Proteome Discoverer 3.3. SEQUEST-HT and CHIMERYS search algorithms were used to match HCD spectra acquired in the Orbitrap and CID ion trap; SEQUEST-HT was used with variable phosphorylation (serine, threonine, and tyrosine) and acetylation (lysine and N-terminus). Two CHIMERYS search nodes were used to allow variable phosphorylation (serine, threonine, and tyrosine) or acetylation (lysine). The same normalization and statistical tests as described above were used. Protein groups with a q-value <0.05 and an absolute log₂ fold difference >0.485 (corresponding to a >1.4-fold difference) were considered differentially abundant.

#### 2.7.4 Additional Data Analyses

**MS2 coverage analysis**: For each LFQ mass feature, all MS2 spectra within 30 ppm precursor mass and 2 min retention time were identified from FLASHDeconv-generated msAlign files. Features were classified as: (i) identified, if a matching TopPIC identification passed the FDR; (ii) fragmented (but unidentified), if MS2 spectra were present but no proteoform was identified; or (iii) not fragmented, if no MS2 spectrum was acquired. **Condition-specific mass features**: Features detected in all replicates of one condition and absent from all replicates of the other condition were classified as condition-specific. **Fragment ion counting**: Fragment ion counts per MS2 spectrum were extracted from msAlign files. For features with multiple MS2 spectra, the spectrum with the highest spectral quality score was used for fragment ion counting. **MS1 sampling density**: The number of MS1 spectra contributing to each mass feature was obtained from the FLASHDeconv mass feature output (column “MassCount”). **Coefficient of variation**: For each mass feature quantified in all replicates, the coefficient of variation (CV) of abundance values was calculated per biological condition as the standard deviation divided by the mean across replicates. **Comparison of MS1-only and MS1/MS2 datasets**: Mass features were matched by precursor mass (20 ppm) and retention time (120 s).

## 3 Results & Discussion

### 3.1 Challenges in Conventional DDA-based LFQ Workflows in TDP

First, a conventional MS1/MS2 experiment was performed to examine factors limiting current TDP-LFQ workflows. For this, the proteomes of *Escherichia coli* grown in minimal medium supplemented with glucose and acetate, respectively, were analyzed, with four biological replicates per condition. After enriching proteoforms suitable for TDP by solid-phase extraction, each sample was analyzed individually by LC-MS/MS using a DDA method. Only mass features detected across all replicates in both conditions (hereinafter referred to as quantified mass features) were included in downstream analysis. Subsequently, identified proteoforms were linked to quantified mass features based on their monoisotopic mass and retention time.

The vast majority of mass features were exclusively detected in a single replicate, and only 434 (5%) were detected across all eight samples (**Figure 2a**), demonstrating a high rate of missing values. Furthermore, of the 873 proteoforms identified, only 220 (25%) could be assigned to mass features that fulfilled the quantitative filtering criteria (**Figure 2b**).

To determine why only a small fraction of quantified mass features could be assigned to proteoforms, we systematically investigated the limiting factors. More than 37% of proteoforms were assigned to mass features that failed the quantitative filtering criteria (i.e., that were not consistently detected across all replicates; **Figure 2b**). Note that the applied mass and retention time tolerances for matching identified proteoforms with quantified mass features were confirmed to be appropriate (**Supplementary Figure 4a**). Furthermore, for 38% of the proteoforms, no corresponding mass feature was detected at all. This is possible since proteoform identification only requires (a single) precursor deconvolution, whereas mass feature detection additionally requires reconstruction of the chromatographic elution profile. Notably, the proteoforms not matching any mass feature showed lower overall identification quality, as reflected in lower E-value distributions and a higher proportion of proteoforms carrying unknown mass shifts (**Supplementary Figure 4b,c**). In addition, they were predominantly observed in the low-molecular-weight range (<5 kDa) and at late retention times (**Figure 2c**). Conversely, proteoforms associated with mass features detected in a larger number of replicates generally exhibited higher identification quality (**Supplementary Figure 4d,e**). Together, these observations suggest that a substantial fraction of proteoforms without corresponding mass features may originate from false-positive identifications, e.g., arising from errors in MS1 deconvolution.^27,28^

On the other hand, only 51% of the 434 quantified mass features were assigned to proteoforms. Approximately 10% of quantified mass features were not selected for MS2 fragmentation during DDA acquisition and therefore remained inaccessible for identification. Moreover, more than 40% of the mass features were fragmented but failed to yield a proteoform identification. Compared with identified proteoforms, those generated substantially fewer fragment ions (median: 13 vs. 46; **Supplementary Figure 4f**), indicating that insufficient fragmentation is a major factor limiting confident proteoform identification. Importantly, this does not necessarily imply suboptimal MS2 acquisition settings, since in DDA-based workflows, MS2 acquisition is inherently constrained by a trade-off between spectral quality and analytical depth: increasing quality (e.g., due to higher resolution, longer injection times, or multiple fragmentation events per precursor) can improve identification quality, but simultaneously reduces the number of precursors that can be analyzed.

Moreover, highly abundant mass features are substantially more likely to be selected for MS2 fragmentation and, once fragmented, more likely to yield proteoform identifications (**Figure 2d**). In contrast, low-intensity mass features showed a markedly reduced probability of both MS2 selection and successful identification, demonstrating the well-known fundamental sampling bias of DDA-based workflows.^29^ For example, approximately 29% of mass features in the lowest-intensity quartile were not selected for fragmentation, whereas only 1% of those in the highest-intensity quartile remained unfragmented.

Differential abundance was assessed using Welch’s t-tests, with multiple-testing correction by the Benjamini-Hochberg procedure. Of the 439 quantified mass features, 49 were significantly differentially abundant between the two conditions (FDR <5%, absolute log_2_ fold-change >0.58). However, only 35 of those could be assigned to identified proteoforms.

In summary, the results reveal three major limitations of current DDA-based TDP-LFQ workflows: (1) high numbers of missing values due to limited reproducibility of mass feature detection; (2) many quantified mass features fail to generate sufficiently informative MS2 spectra; (3) low-abundance mass features are preferentially undersampled. Together, these factors substantially reduce the overlap between quantified mass features and identified proteoforms, thereby limiting both the analytical depth and biological interpretability of current TDP-LFQ experiments.

### 3.2 MS1-only Approach for Improved Label-free Quantification

Since missing values were identified as a major limitation of conventional TDP-LFQ workflows, we next investigated an MS1-only acquisition strategy for proteoform quantification. By omitting MS2 scans, the entire measurement time is available for MS1 mass feature detection, thereby increasing the number of MS1 spectra acquired across chromatographic elution profiles. We hypothesized that this increased sampling density would improve the reproducibility of mass feature detection across replicates and, consequently, reduce the number of missing values.

MS1-only measurements were acquired for the same samples analyzed above using identical MS1 acquisition settings as in the MS1/MS2 approach. Overall, the MS1-only approach detected 18% more mass features than the MS1/MS2 workflow (**Figure 3a**) and, more importantly, the number of mass features quantified across all replicates of both conditions increased by 162%. The overlap of quantified mass features was very high between the two approaches: 743 were exclusively quantified in the MS1-only approach, 396 in both approaches, and only 38 exclusively in the conventional MS1/MS2 workflow (**Supplementary Figure 5a**). Moreover, in the MS1/MS2 workflow, 66% of all detected features were observed in only a single replicate, whereas this number decreased to 45% in the MS1-only approach (**Figure 2a, Supplementary Figure 6**). In addition, the number of condition-specific mass features (i.e., mass features consistently detected in all replicates of one condition while being absent in all replicates of the other condition) increased by 230% under acetate (152 vs. 46) and 130% under glucose (85 vs. 37) conditions using the MS1-only approach compared with the conventional MS1/MS2 workflow.

**Figure 3:**
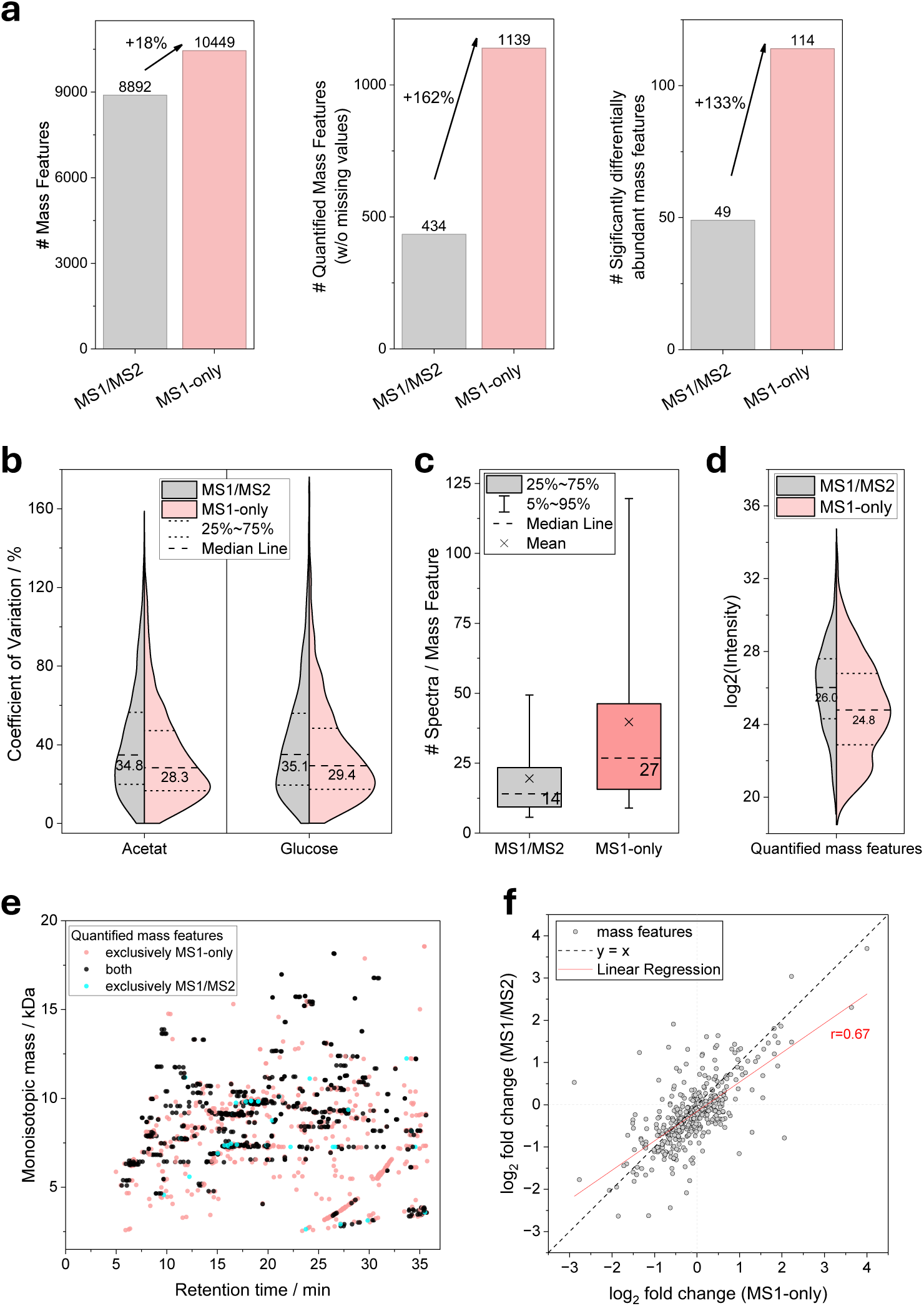
Comparison of conventional MS1/MS2 and MS1-only workflows for mass feature quantification. **a** Number of detected (left; detected in at least one replicate), quantified (middle; detected in all replicates of both conditions), and significantly differentially abundant (right; Welch’s t-test, Benjamini–Hochberg correction, FDR <1%, fold change >1.5) mass features. **b** Distribution of coefficients of variation (scale: count). **c** Distribution of the number of MS1 spectra per quantified mass feature. **d** Distribution of mass feature intensities (scale: count). **e** Scatter plot of quantified mass features by precursor mass and retention time. **f** Comparison of log₂ fold changes obtained with the MS1-only and conventional MS1/MS2 workflows. Each point represents a mass feature quantified by both workflows. The dashed line indicates identical fold changes (y=x), and the red line shows the linear regression.

Pearson correlations of mass feature intensities showed high reproducibility between biological replicates within each condition, while clearly separating the two cultivation conditions (**Supplementary Figure 7a,b)**. Compared with the conventional MS1/MS2 workflow, the MS1-only approach consistently yielded slightly higher replicate correlation coefficients. Furthermore, the coefficient of variation (CV) was used as a measure of quantification precision (**Figure 3b**). For the MS1/MS2 approach, median CV values for glucose and acetate replicates were approximately 35%. In contrast, substantially lower CV values were observed for the MS1-only approach, with median CVs of 29% and 28%. The improved precision observed with the MS1-only quantification can be attributed to the substantially increased number of MS1 spectra acquired per mass feature (median: 27 vs. 14, **Figure 3c**). The higher spectral sampling density enables more accurate reconstruction of chromatographic elution profiles and reduces stochastic fluctuations between replicate measurements.

Furthermore, the MS1-only approach additionally quantified mostly low-abundance mass features (**Figure 3d**) that covered the entire mass and retention time range (**Figure 3e**), highlighting that the improved sensitivity was not restricted to a specific subset of proteoforms. For both acquisition strategies, detection reproducibility strongly depends on signal intensity: highly abundant mass features were more consistently observed across replicates, whereas low-abundance features were more prone to missing values (**Supplementary Figure 7c,d)**. A small number of mass features represented signals from common polymer contaminants (such as polyethylene glycol), predominantly eluting at the end of the gradient (**Supplementary Figure 7e,f**). This observation highlights an inherent limitation of label-free MS1-based quantification because non-proteoform signals cannot be unambiguously distinguished without MS2 information.

After statistical analysis of the 1,139 quantified mass features detected across all replicates in the MS1-only approach, 114 were classified as significantly differentially abundant (FDR <5%, absolute log_2_ fold-change >0.58), representing an increase of +133% compared to the conventional MS1/MS2 approach (**Supplementary Figure 8**).

The overall correlation of fold changes for mass features quantified in both acquisition strategies was high (Pearson correlation: 0.67), showing that both workflows captured consistent quantitative trends (**Figure 3f**). Among the significantly differentially abundant mass features, most (n=89) were unique to the MS1-only workflow, whereas 25 were shared between the workflows, and 24 were unique to the conventional MS1/MS2 workflow (**Supplementary Figure 5b**). Almost all mass features that reached statistical significance in only one workflow showed comparable fold changes in the other dataset (**Supplementary Figure 9**), indicating that the observed differences were primarily driven by statistical confidence rather than discordant quantitative data. The increased number of significantly differentially abundant mass features detected by the MS1-only approach likely reflects substantially improved quantitative precision from increased MS1 sampling density. This reduced replicate variability thereby increases statistical power. A small subset of mass features reached statistical significance exclusively in the conventional MS1/MS2 workflow, possibly reflecting the dataset’s lower analytical complexity, where reduced sensitivity yields fewer co-eluting low-abundance mass features and, consequently, less signal interference. These effects could improve statistical power for a small subset of features, despite the overall reduction in quantitative data quality in the MS1/MS2 approach.

To evaluate the robustness and broad applicability of the MS1-only workflow, the approach was further assessed across different workflows and experimental designs. Increasing the gradient length from 30 min to 60 min nearly doubled the number of quantified mass features in both acquisition strategies (**Supplementary Figure 10a**). The MS1-only workflow quantified substantially more mass features (+172%) than the corresponding MS1/MS2 measurements (MS1/MS2: 826; MS1-only: n=2,243), resulting in 97% more differentially abundant mass features (116 vs. 229). The same samples, analyzed using 90-min gradients without FAIMS, yielded 1,355 and 2,878 quantified mass features using the MS1/MS2 and MS1-only approaches, with 64 and 101 differentially abundant mass features (+58% more in the MS1-only approach) (**Supplementary Figure 10b**).

Furthermore, an additional dataset was acquired in which the *Escherichia coli* proteoforms were reduced, alkylated, purified, and analyzed without prior enrichment of small proteoforms. Using a 30-min gradient on an MAbPac column without utilizing FAIMS, 130 and 335 mass features were quantified using the conventional MS1/MS2 and the MS1-only workflows, resulting in the identification of 10 and 41 significantly differentially abundant mass features (i.e., a 310% increase for the MS1-only approach; **Supplementary Figure 10c**). Notably, the lower number of significantly differentially abundant mass features obtained without FAIMS demonstrates that FAIMS also improves proteoform-level quantification, consistent with previous findings by Kline et al.^9^

In summary, the MS1-only workflow substantially reduced missing values, improved quantitative precision, and increased the number of quantified and significantly differentially abundant mass features. The superiority of the MS1-only strategy was reproducibly observed across different gradient lengths, sample preparation workflows, and acquisition setups, demonstrating its broad applicability and robustness.

### 3.3 Targeted Data-dependent MS2 Acquisition for Proteoform Identification

Since the MS1-only approach exclusively acquires quantitative mass feature information, separate LC-MS/MS runs are required for proteoform identification. In a first experiment, we targeted significantly differentially abundant mass features (determined in the MS1-only measurements) using the same MS2 acquisition settings as in the conventional MS1/MS2 approach. If no target mass was identified in an acquired MS1 spectrum, the most abundant precursors were selected for fragmentation. Across three replicates, 327 proteoforms were identified, including 51 proteoforms that were significantly differentially abundant. Compared with the conventional MS1/MS2 workflow, this corresponded to a 46% increase in the number of differentially abundant proteoforms.

Although targeted data-dependent MS2 acquisition substantially increased the number of differentially abundant proteoforms, only 44% of the targeted mass features could be identified. Consistent with the findings from the conventional MS1/MS2 workflow, proteoform identification remained primarily limited by insufficient fragment ion generation and the low abundance of precursor mass features (**Supplementary Figure 11**).

To address these limitations, the targeted MS2 workflow was further optimized to improve both MS2 sensitivity and fragment ion generation: MS2 spectra were acquired at higher resolution (120k) with longer maximum injection times (246 ms). In addition, two fragmentation spectra were acquired per precursor using HCD (30% NCE) and CID (35% NCE). The rationale for this approach was that different proteoforms may fragment more efficiently under different fragmentation conditions, thereby increasing identification depth. Using these optimized MS2 settings, across three replicates, 244 proteoforms were identified, including 67 that were significantly differentially abundant (**Figure 4**). Compared with the initial targeted MS2 acquisition, the optimized workflow substantially reduced the number of fragmented but unidentified precursors (**Supplementary Figure 12a,b**). Furthermore, the improved MS2 sensitivity enabled the identification of additional low-abundance proteoforms and significantly increased the number of detected fragment ions per proteoform (**Supplementary Figure 12c,d**).

**Figure 4:**
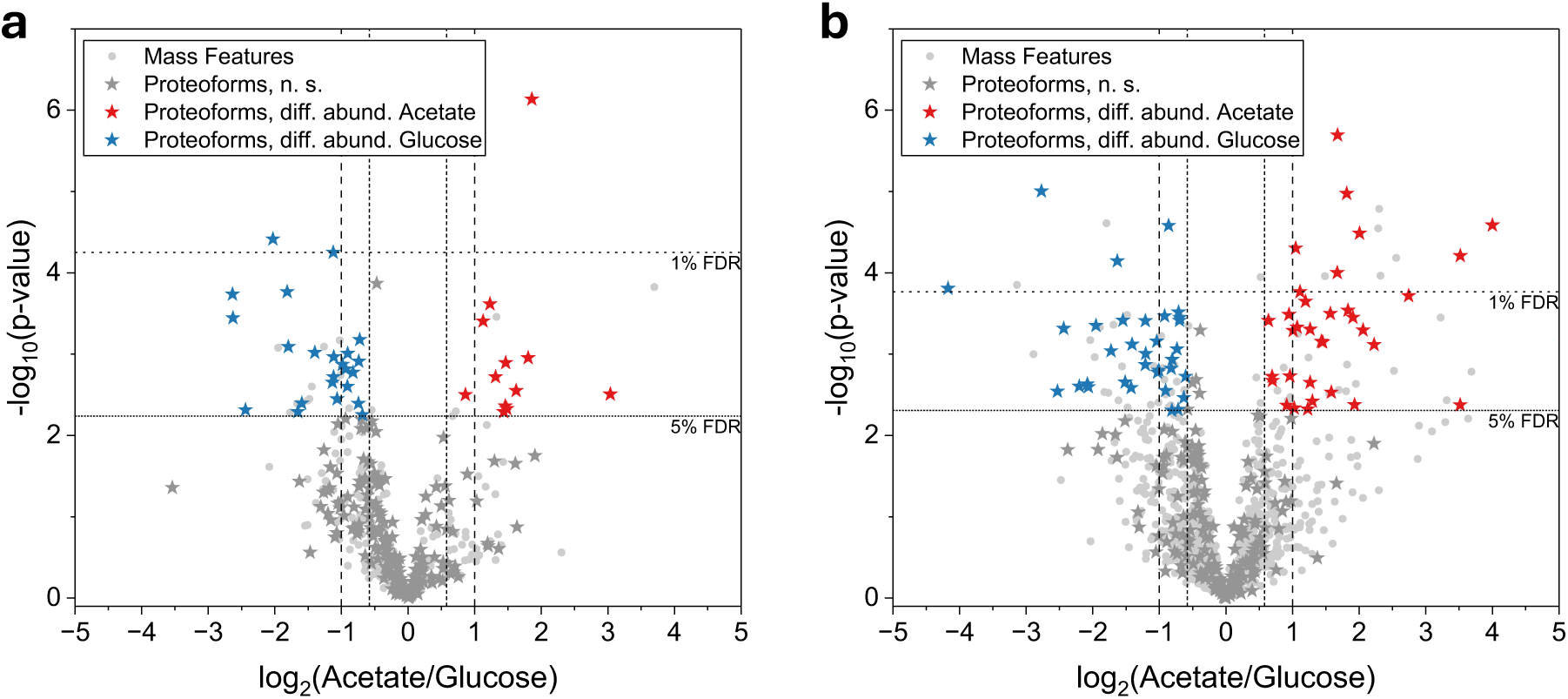
Comparison of quantified proteoforms obtained with the conventional MS1/MS2 workflow and the developed MS1-only/targeted MS2 workflow. **a** Conventional MS1/MS2 workflow. **b** MS1-only/targeted MS2 workflow. Differentially abundant mass features were targeted, and two high-quality fragmentation spectra were acquired. Statistical significance at the mass feature level was assessed using Welch’s t-test with Benjamini–Hochberg correction. The stars represent the proteoforms, and the dots represent mass features that do not result in proteoform identifications. Colored stars indicate significantly (FDR <5%) differentially abundant proteoforms with an absolute log_2_ fold change >0.58 (red, higher abundance in acetate; blue, higher abundance in glucose). Horizontal lines: 5% and 1% FDR cutoffs; vertical lines: log_2_ fold changes of ±0.58 and ±1.

Next, all quantified mass features were targeted using the optimized MS2 settings to maximize proteome coverage. Using the optimized acquisition parameters (120k resolution, 246 ms maximum injection time, and two MS2 spectra per precursor), a total of 326 proteoforms were identified across all three replicates, of which 60 were significantly differentially abundant (**Supplementary Figure 13**). Compared with the 220 quantified proteoforms obtained with the conventional MS1/MS2 workflow, this corresponded to an increase of 48%. However, a substantial fraction of low-abundance precursors remained unselected for fragmentation. This likely resulted from the increased MS2 injection times and the acquisition of multiple MS2 scans per precursor, which reduced the number of distinct precursors that could be fragmented during chromatographic elution. In addition, some proteoforms likely remained difficult to identify due to intrinsically poor fragmentation behavior or insufficient fragment ion generation under the applied conditions. Furthermore, a subset of the targeted mass features may not represent genuine proteoforms but instead originate from contaminating species, such as polymer contaminants (see above).

The targeted MS2 strategy also enabled selective analysis of condition-specific proteoforms. Whereas the MS1/MS2 approach identified 14 and 6 proteoforms exclusively detected under acetate and glucose conditions, respectively, the MS1-only/targeted MS2 workflow identified 19 (+36%) and 16 (+167%) condition-exclusive proteoforms. The comparatively low number of identified condition-exclusive proteoforms relative to the corresponding quantified mass features reflects the challenges of MS2-based identification of low-abundance proteoforms.

To further validate the MS1-only/targeted MS2 workflow, the experiment was repeated using the same samples and analytical setup but with a 60 min chromatographic gradient. In the conventional MS1/MS2 approach, this resulted in 432 quantified proteoforms, of which 84 were significantly differentially abundant (**Supplementary Figure 14**). Following MS1-only acquisition and targeted MS2 analysis of the significantly differentially abundant mass features (triplicates), a total of 522 proteoforms (+21%) were identified across three replicates, including 118 significantly differentially abundant proteoforms (+40%). These results demonstrate that the benefit of MS1-only/targeted MS2 acquisition is not restricted to a specific chromatographic setup but is maintained under conditions that increase proteoform coverage by extending the gradient length.

In summary, combining MS1-only quantification with targeted MS2 acquisition substantially increases both proteoform quantification depth and the number of proteoforms that are significantly differentially abundant, compared with conventional MS1/MS2 workflows. Importantly, decoupling quantification and identification enables independent optimization of both acquisition strategies and allows MS2 acquisition time to focus specifically on quantitatively relevant mass features.

### 3.4 Biological Insights from Proteoform-Level Quantification

To demonstrate the biological utility of the proposed workflow, proteoform abundance changes were analyzed in *Escherichia coli* grown on minimal medium supplemented with glucose or acetate as the sole carbon source. Quantification was based on 60-min MS1-only measurements, and proteoform identifications were obtained by combining all acquired targeted and untargeted MS2 acquisitions. After filtering to mass features detected in all four biological replicates of at least one condition, we imputed missing values from a down-shifted normal distribution and applied the same statistical testing as described before, resulting in 4,424 quantified mass features. TopPIC database search allowed the assignment of 1,392 proteoforms to those quantified mass features, of which 315 showed a significant difference in abundance between the two growth conditions (**Figure 5a; Supplementary Table 1**). The proteoform-level dataset obtained by TDP was then compared with a previously acquired^15^ and reanalyzed comprehensive 2D-LC-MS BUP isobaric labeling-based quantitative dataset, with 2,283 protein-level quantifications (**Supplementary Figure 15; Supplementary Table 2**).

**Figure 5:**
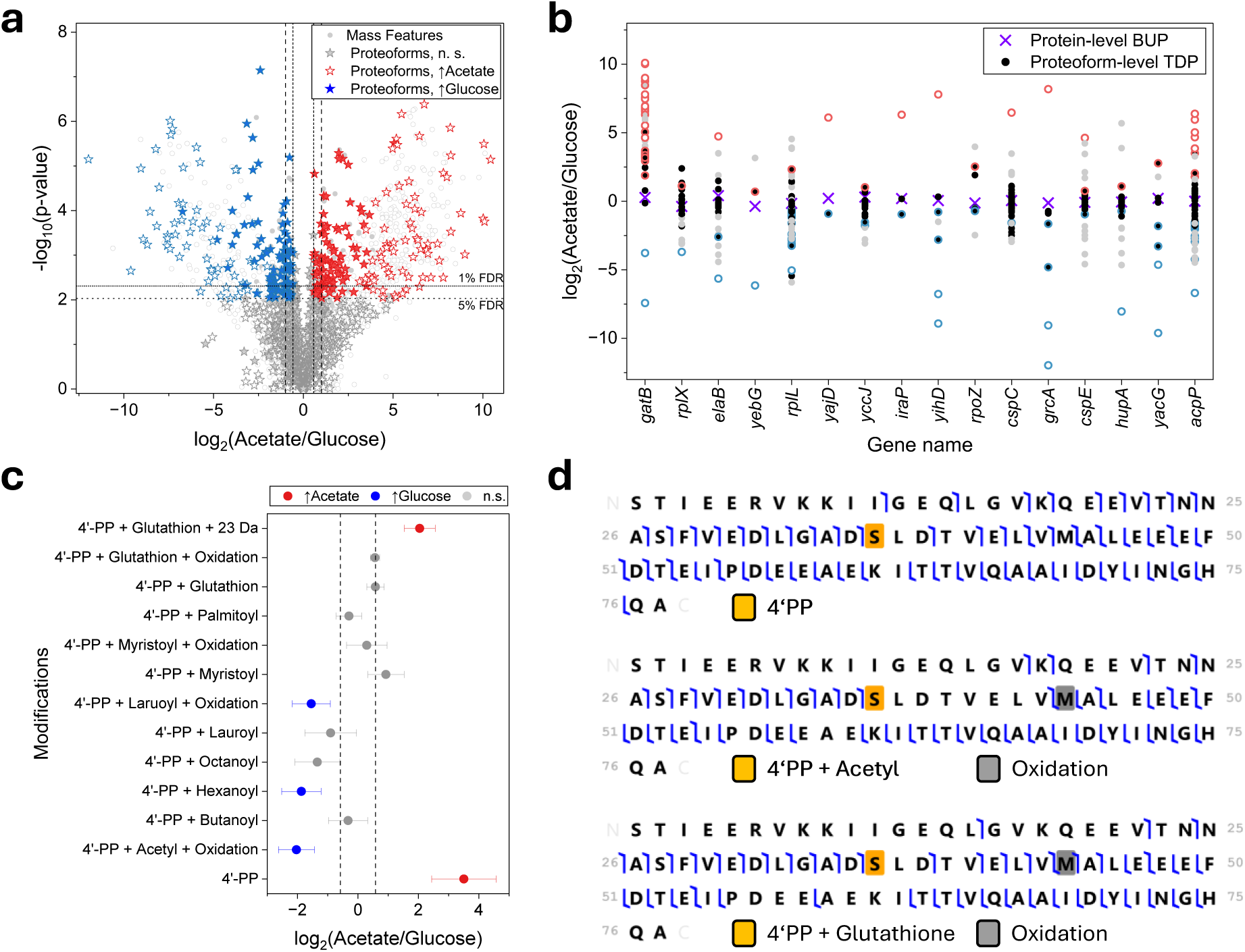
High analysis depth revealed biological insights not accessible with bottom-up proteomics. Statistical analysis was based on MS1-only acquisitions using a 60-min gradient and multiple targeted and untargeted MS2 acquisitions. Mass features that were detected in all four biological replicates of at least one condition were used for quantification, and data imputation for missing values was applied. **a** Volcano plot of quantified mass features and assigned proteoforms. Unfilled stars/dots indicate quantified data obtained only after imputation. Horizontal lines: 5% and 1% FDR cutoffs; vertical lines: log_2_ fold changes of ±0.58 and ±1. **b** Comparison of proteoform-level TDP with protein-level BUP quantifications. Shown are proteins that were quantified by BUP and TDP, with at least one significantly differentially abundant proteoform under both conditions in the TDP dataset. Dots represent individual proteoforms from a given protein quantified by TDP, and crosses represent protein-level data quantification obtained by BUP (reanalysis from a previous. Colored dots indicate significantly differentially abundant proteoforms, and unfilled dots represent imputed data. **c** Selected quantified proteoforms of the Acyl Carrier Protein (AcpP, UniProt accession: B1XA04; without data imputation). 4’PP, phosphopantetheine. **d** Fragment maps of the Acyl Carrier Protein proteoform with O-(pantetheine 4’-phosphoryl)serine (4’-PP, upper panel; ΔM=−2.40 ppm; p-value 3.2e-107, C-score 1,293, 72% residue cleavage), 4’-PP with a covalently bound aetcyl group and methionine oxidation (middle panel; ΔM=−1.94 ppm, p-value 1.7e-80, C-score 942, 55% residue cleavage), and 4’-PP with a covalently bound glutathione and methionine oxidation (lower panel; ΔM=1.88 ppm, p-value 7e-86, C-score 1,013, 58% residue cleavage).

In the TDP dataset, 166 of 237 (70%) proteins were represented by multiple quantified proteoforms (**Supplementary Figure 16a**). Among those, 96 proteins had at least one significantly differentially abundant proteoform. Notably, for 17 proteins, individual proteoforms showed opposing abundance changes between the two growth conditions, with at least one proteoform being more abundant in each condition. In contrast, BUP provided quantitative data for 16 of these proteins, none of which showed significant protein-level differences between conditions (**Figure 5b**). This highlights that protein-level quantification based on summed peptide-level data can obscure biologically relevant differences in proteoform abundance. Such differences may be missed due to limitations in the experimental setup (e.g., proteolytic digestion or database search space constraints) or masked when proteoforms with opposing abundance changes are aggregated, rendering the overall protein-level change insignificant (**Supplementary Figure 16b)**.

TDP can reveal proteoforms with different disulfide-bond states, which BUP typically cannot resolve. For example, a proteoform of the DNA gyrase inhibitor YacG (UniProt accession B1XC77) containing two disulfide bridges was of higher abundance under growth on glucose (log_2_ fold change −1.8). In contrast, the proteoform with free sulfhydryl groups was more abundant under acetate growth conditions (log_2_ fold change +2.78) (**Supplementary Figure 17**).

The bottom-up proteome analysis showed that fatty acid metabolism is among the pathways significantly affected by the different carbon sources. While proteins involved in beta-oxidation were more abundant on acetate growth (mean log2 fold change +0.47), proteins involved in fatty acid synthesis were generally not significantly changed (mean log2 fold change −0.19). In contrast, TDP revealed substantial condition-dependent remodeling at the proteoform level. A notable example is the acyl carrier protein (AcpP; UniProt accession B1XA04), which carries acyl intermediates through its 4′-phosphopantetheine (4′-PP) prosthetic group. AcpP was represented by dozens of quantified proteoforms in the TDP dataset (**Figure 5c**). While at the protein level, no change in AcpP abundance could be detected, its proteoform composition differed significantly between the two conditions, with distinct molecular states preferentially abundant under glucose or acetate growth, respectively. The holo-AcpP and an additional proteoform with the active 4′-PP site protected by glutathione modification were more abundant under acetate growth conditions, whereas several different acylated 4′-PP proteoforms directly involved in subsequent elongation cycles of fatty acid synthesis were more abundant under growth on glucose (**Figure 5c,d**).

## 4 Conclusion

Based on a systematic investigation of the limitations of conventional DDA-based TDP-LFQ, we developed a dual-step TDP-LFQ workflow consisting of initial MS1-only acquisitions for sensitive mass feature quantification, followed by targeted data-dependent MS2 measurements for proteoform identification. The feasibility of performing MS1-only quantification and subsequent MS2-based proteoform identification separately has previously been demonstrated by Durbin et al.^12^ By decoupling mass feature quantification and proteoform identification, both acquisitions can be independently optimized without negatively affecting each other.

Conversely, conventional LFQ workflows combine MS1-based quantification and DDA-based MS2 identification within a single acquisition, creating an inherent trade-off: MS1 scans are required for accurate, reproducible quantification, while MS2 scans are required for proteoform identification. Increasing the time spent on MS1 or MS2 levels reduces the time available for the other.

MS1-only acquisition nearly doubled the number of quantified mass features and substantially improved quantitative precision. Targeted MS2 measurements enabled selective fragmentation of relevant mass features and can be fully tailored and optimized (e.g., based on proteoform mass ranges)^30^ to maximize proteoform identification depth. The advantages of the developed workflow were consistently reproduced across different chromatographic gradients, sample preparation workflows, and instrumental setups, highlighting its broad applicability. A conceptually related strategy using MS1-only acquisition has been established in BUP through accurate mass and time tag approaches.^31^

Notably, the MS1-only and targeted MS2 workflow requires longer overall acquisition times than conventional MS1/MS2 measurements. In the present study, the conventional workflow required eight LC-MS/MS measurements (four biological replicates per condition), whereas the MS1-only/targeted MS2 workflow required eight MS1-only runs in addition to three targeted MS2 measurements for proteoform identification. Nevertheless, the moderate increase in acquisition time translated into a substantial increase in proteoform coverage and biological information. In addition, the increased sensitivity of MS1-only acquisition also increases the detection of non-proteoform mass features originating from contaminants or chemical background signals, which cannot be unambiguously distinguished in the absence of MS2 information. However, known contaminant-associated mass features can easily be excluded from downstream quantitative analyses.

The results of this study also highlight several general challenges in TDP, including limited reproducibility and the lack of established quality-control metrics for deconvolved mass features, as well as the need for orthogonal validation strategies. A common deconvolution artifact is isotopic errors, which occur when incorrect isotopes are assigned as the monoisotopic peak. Notably, after feature alignment, potential isotopic errors accounted for a similar proportion of mass features in both the conventional MS1/MS2 and MS1-only datasets. Manually excluding these features reduced the total number of features by approximately 15%, while clearly preserving the improvements achieved with MS1-only compared to the conventional MS1/MS2 acquisition (**Supplementary Figure 18**). Thus, the presence of isotopic errors does not affect the conclusions of this study. However, an important future task will be algorithmically identifying and merging isotopic errors directly at the mass feature level, which reduces the number of features subjected to statistical testing and potentially further improves the statistical power of TDP-LFQ analyses. Moreover, since deconvolved masses form the foundation for proteoform identification, the reliability of subsequent data analysis critically depends on the accuracy and reproducibility of MS1-level deconvolution. This is particularly true for database searches utilizing wide precursor tolerances, as unknown mass shifts are calculated as the difference between the theoretical proteoform mass and the experimentally determined mass. Therefore, FDR estimation approaches for deconvolved MS1 masses are required for improving the reproducibility and reliability of both proteoform quantification and identification.^32,33^ Furthermore, retention or migration time information from LC-MS or CE-MS,^34^ or techniques providing information such as lysine-/ or cysteine-count measurements,^33,35^ can provide complementary evidence to increase confidence in proteoform assignments.

Future developments may further refine the presented workflow, including the use of FAIMS with internal compensation voltage stepping,^36^ optimization of MS/MS fragmentation strategies, and intelligent acquisition strategies for automation.^37^ In the future, proteoform atlas-based identification approaches^38,39^ solely based on intact masses and retention/migration times^24^ would diminish the need for targeted MS2 acquisition, further improving analysis throughput.

In summary, the presented workflow demonstrates that decoupling quantitative mass feature analysis from proteoform identification substantially improves quantitative depth and proteoform coverage in TDP-LFQ, providing a flexible platform for high-sensitivity proteoform analysis. Furthermore, the comparison with BUP emphasizes the importance of proteoform-resolved quantification, as protein-level aggregation of peptide measurements can obscure differential abundance and mask biologically relevant proteoform-specific changes.

## Supporting Information

The Supporting Information contains Supplementary Figures 1-17.

## Data Availability

The raw data, deconvolution results, and database search results have been uploaded to the ProteomeXchange Consortium via the PRIDE partner repository with the dataset identifier PXD083443.^26^

## Acknowledgments

The authors thank the German Research Funding organization (DFG) for funding of the Eclipse MS (INST 257-776-1 FUGG).

