## Supporting Information for "Decoupling Quantification and Identification for Label-Free Top-Down Proteomics Improves Proteome Coverage"

### Supplementary Figures

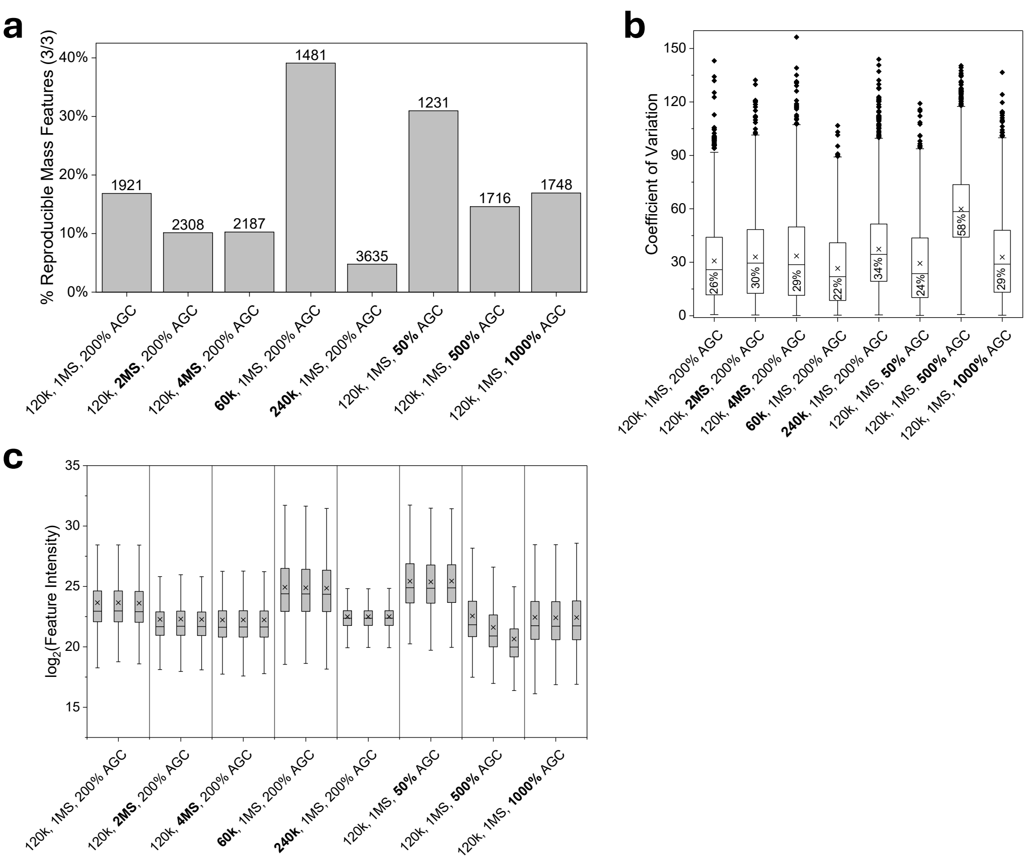


**Supplementary Figure 1**: **Influence of various MS1 settings on mass feature quantification**. Three replicates of *Escherichia coli* proteoforms (full lysate; proteoforms reduced and alkylated) were analyzed at different Orbitrap resolutions, numbers of microscans, and AGC targets (triplicate measurements). The standard settings of the MS1-only methods were: 120k resolution, 1 microscan (MS), 200% AGC target. Mass feature detection and quantification were performed with FLASHDeconv. **a** Percentages and absolute numbers (labels) of mass features reproducibly detected in all three replicates. **b** Coefficient of variation for mass features quantified in all three replicates. **c** Feature intensity of all detected mass features.

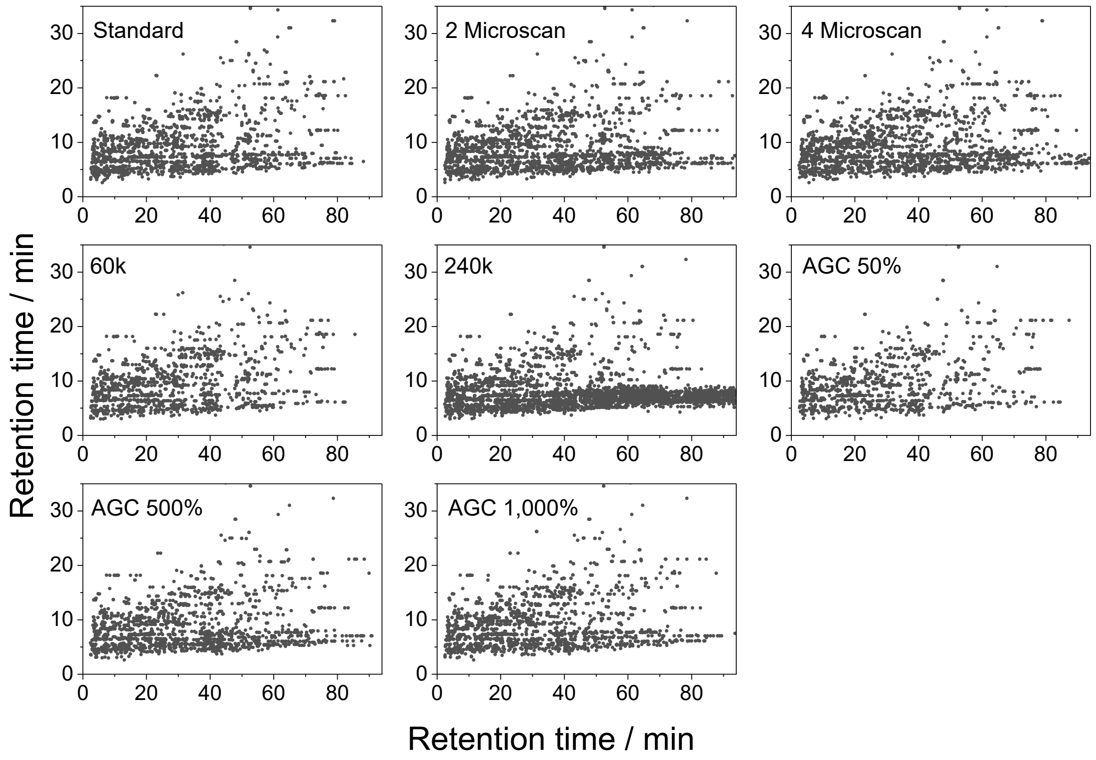


**Supplementary Figure 2**: **Influence of various MS1 settings on mass feature detection**. Three replicates of *Escherichia coli* proteoforms (full lysate; proteoforms reduced and alkylated) were analyzed at different Orbitrap resolutions, numbers of microscans, and AGC targets (triplicate measurements). The standard settings of the MS1-only methods were 120k resolution, 1 microscan (MS), 200% AGC target. Labels indicate the acquisition parameters modified relative to the standard settings.

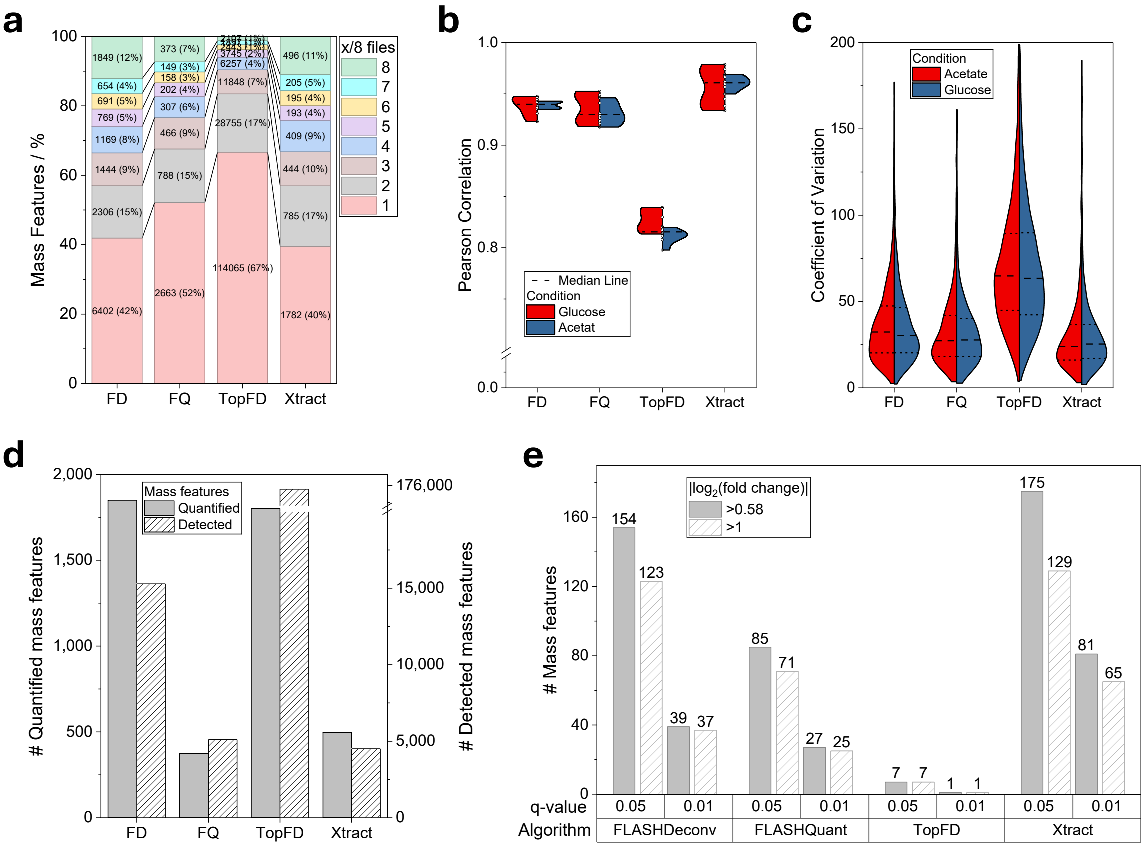


**Supplementary Figure 3**: **Influence of various deconvolution algorithms on mass feature quantification**. Each of the four biological replicates of *Escherichia coli* cultivated in minimal medium supplemented with acetate and glucose, respectively, was analyzed by LC-MS (MS1-only). Deconvolution was performed with FLASHDeconv (FD), FLASHQuant (FQ), TopFD, and Xtract utilizing standard settings. For FLASHDeconv, minimum precursor mass and minimum charge state were set to 2,000 Da and +4, respectively. **a** Percentage of detected mass features detected in certain numbers of raw files. **b** Pearson correlations across all replicates (scale: count). **c** Coefficient of variation for mass features detected in all replicates (scale: width). **d** Number of all detected and quantified (i.e., detected in all eight replicates) mass features. **e** Number of significantly differentially abundant mass features after statistical analysis (Welch's t-test, Benjamini-Hochberg false-discovery rate (FDR)).

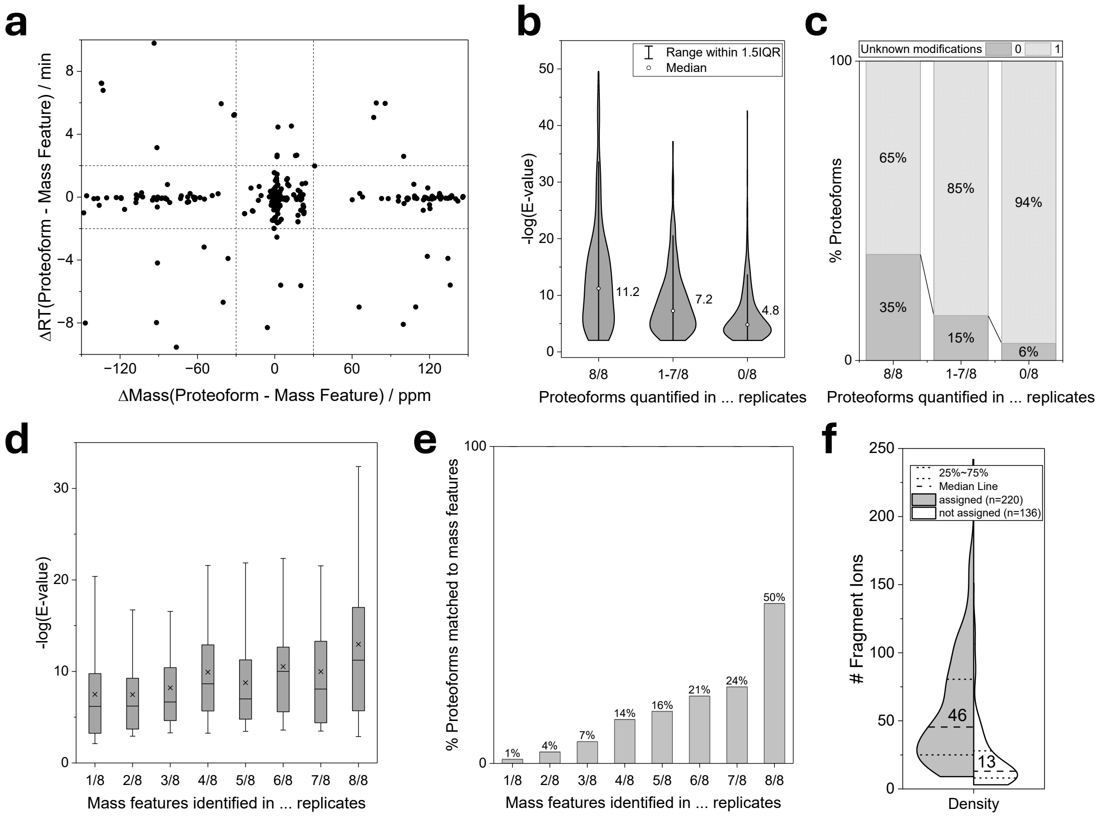


**Supplementary Figure 4**: **Factors limiting the depth of conventional MS1/MS2 DDA-based quantification strategies**. **a** Mass and retention time deviations between identified proteoforms and the closest matching (based on mass) quantified mass features. The dashed lines indicate the matching tolerances (±30 ppm and ±2 min). **b** Distribution of TopPIC E-values for proteoforms assigned to mass features quantified in all eight replicates (8/8), only a subset of replicates (1–7/8), or not detected as mass features (0/8) (scale: count). **c** Fraction of proteoforms carrying unknown mass shifts in the three quantification categories shown in (b). **d** TopPIC E-value distributions as a function of the number of replicates in which the corresponding mass feature was detected. **e** Percentage of detected mass features that could be assigned to proteoform identifications as a function of the number of replicates in which the mass feature was detected. **f** Distribution (scale: count) of fragment ions in MS2 spectra of quantified mass features (in 8/8 replicates) that were either successfully assigned to proteoforms (assigned) or remained unidentified (not assigned).

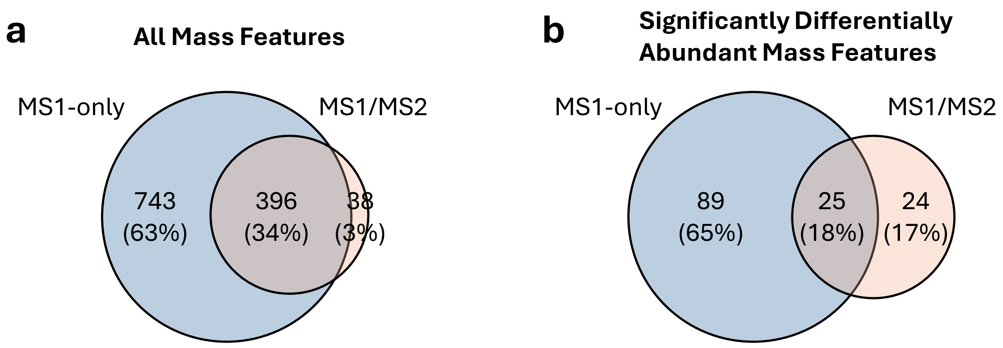


**Supplementary Figure 5**: **Overlap of quantified mass features between the MS1-only and MS1/MS2 approaches**. **a** All quantified (detected in all 8 replicates) mass features. **b** Significantly differentially abundant mass features.

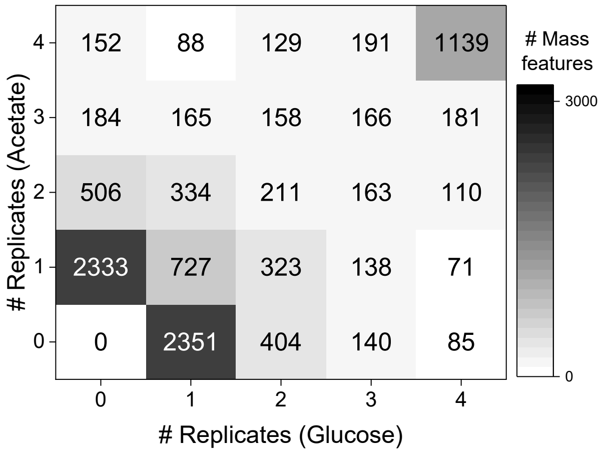


**Supplementary Figure 6**: **Reproducibly detected mass features in the MS1-only approach.** Shown are the numbers of mass features detected across different numbers of biological replicates.

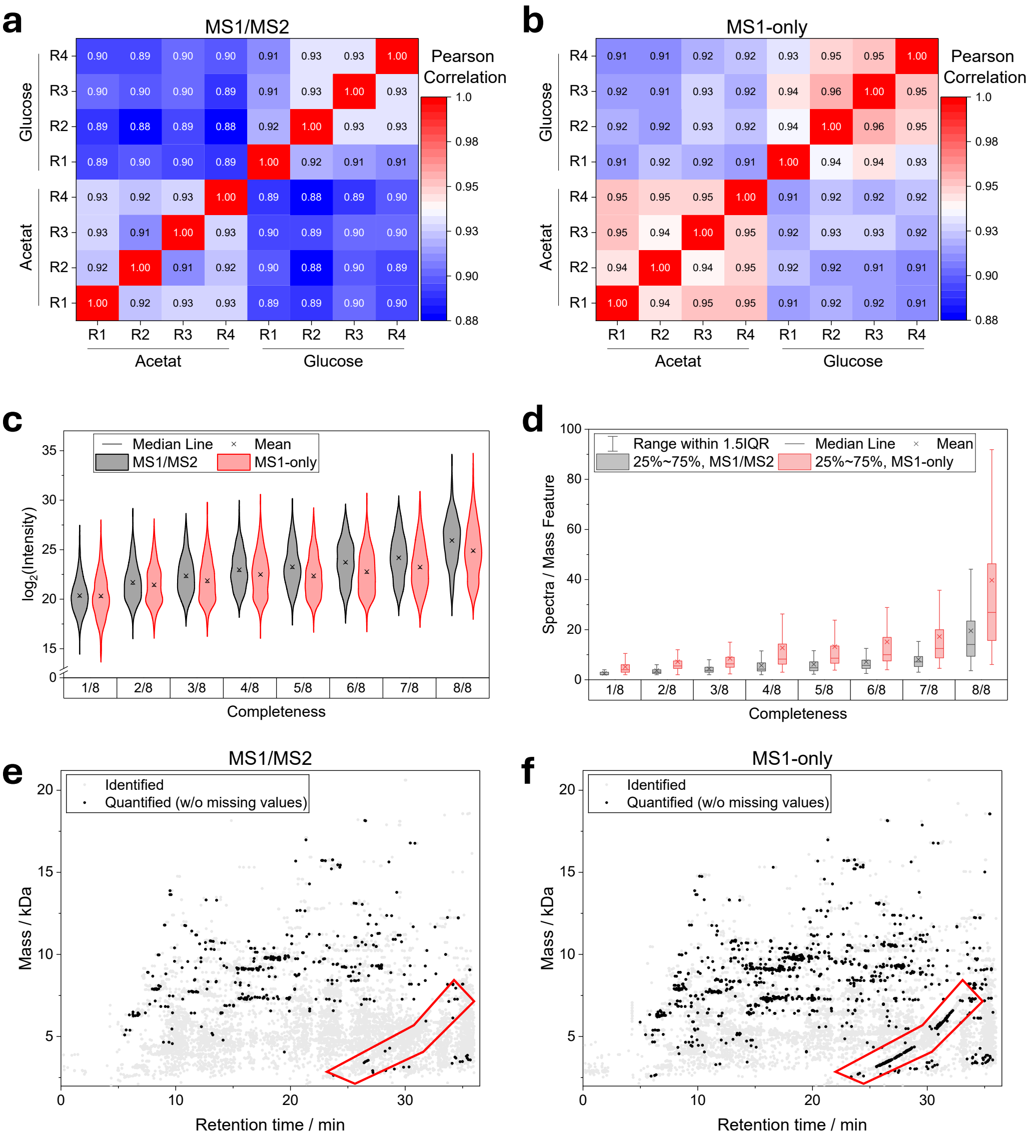


**Supplementary Figure 7**: **Comparison of detected mass features quantified in all replicates between the MS1/MS2 and the MS1-only approaches**. **a,b** Pearson correlation between replicates for the MS1/MS2 and the MS1-only approaches. **c,d** Distributions of (**c**) mass feature intensities (scale: width) and (**d**) spectra per mass feature depending on the number of replicates in which the mass feature was detected. **e,f** Distribution of detected (in 1-7/8 replicates) and quantified (8/8 replicates) mass features regarding retention time and molecular weight. The red boxes indicate common polymer contaminations (such as polyethylene glycol with characteristic MS1 mass shift of ΔM=44 Da).

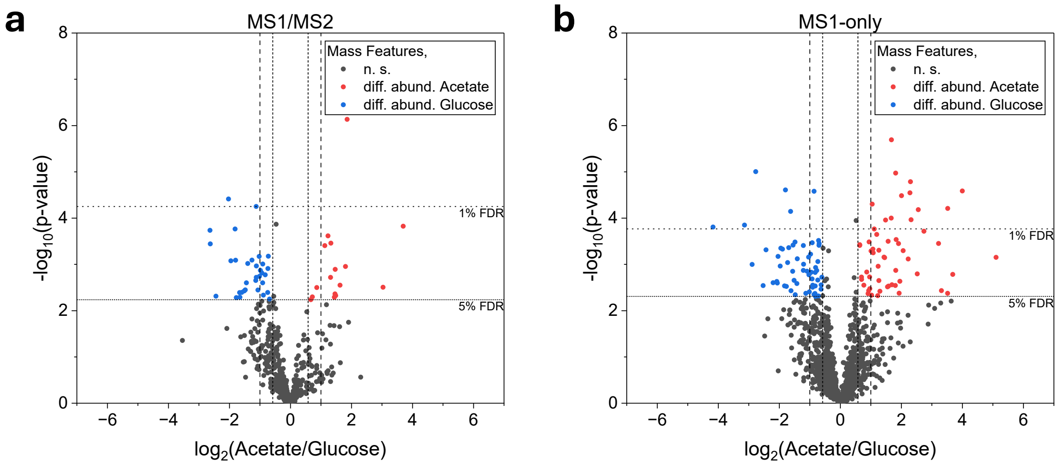


**Supplementary Figure 8**: **Statistical evaluation at the mass feature level**. Shown are volcano plots after analysis using either **a** the conventional MS1/MS2 or **b** the MS1-only approach. Colored dots: significantly differentially abundant mass features more abundant in glucose (blue) or acetate (red) conditions. Horizontal lines: 5% and 1% FDR cutoffs (Benjamini-Hochberg correction); vertical lines: $\log_2$ fold changes of ±0.58 and ±1.

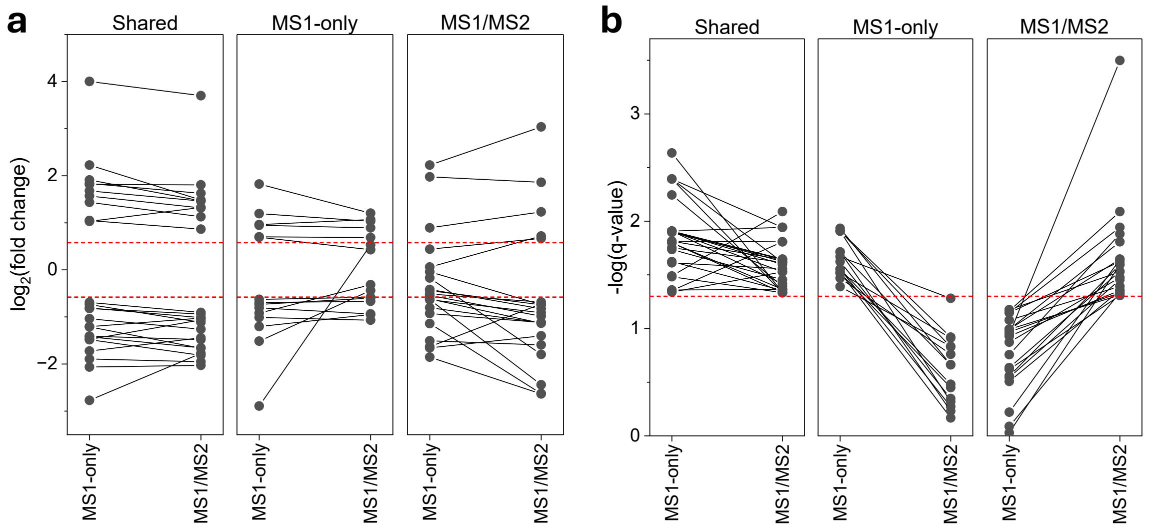


**Supplementary Figure 9**: **Comparison of quantitative results obtained with the MS1-only and conventional MS1/MS2 workflows**. Each point represents a mass feature that was quantified in all four replicates in both biological conditions by both workflows. **a** Comparison of $\log_2$ fold changes determined by the MS1-only and conventional MS1/MS2 workflows. The red dashed line indicates fold change cutoffs (absolute $\log_2$ fold change >0.58). **b** Comparison of −log(q-values) obtained by the two workflows. The red dashed line indicates FDR cutoffs (−log(q) >1.3; FDR <5%). Shared: mass features that were identified as significantly differentially abundant in both workflows. MS1-only: mass features that were identified as significantly differentially abundant exclusively in the MS1-only strategy. MS1/MS2: mass features that were identified as significantly differentially abundant exclusively in the MS1/MS2 strategy.

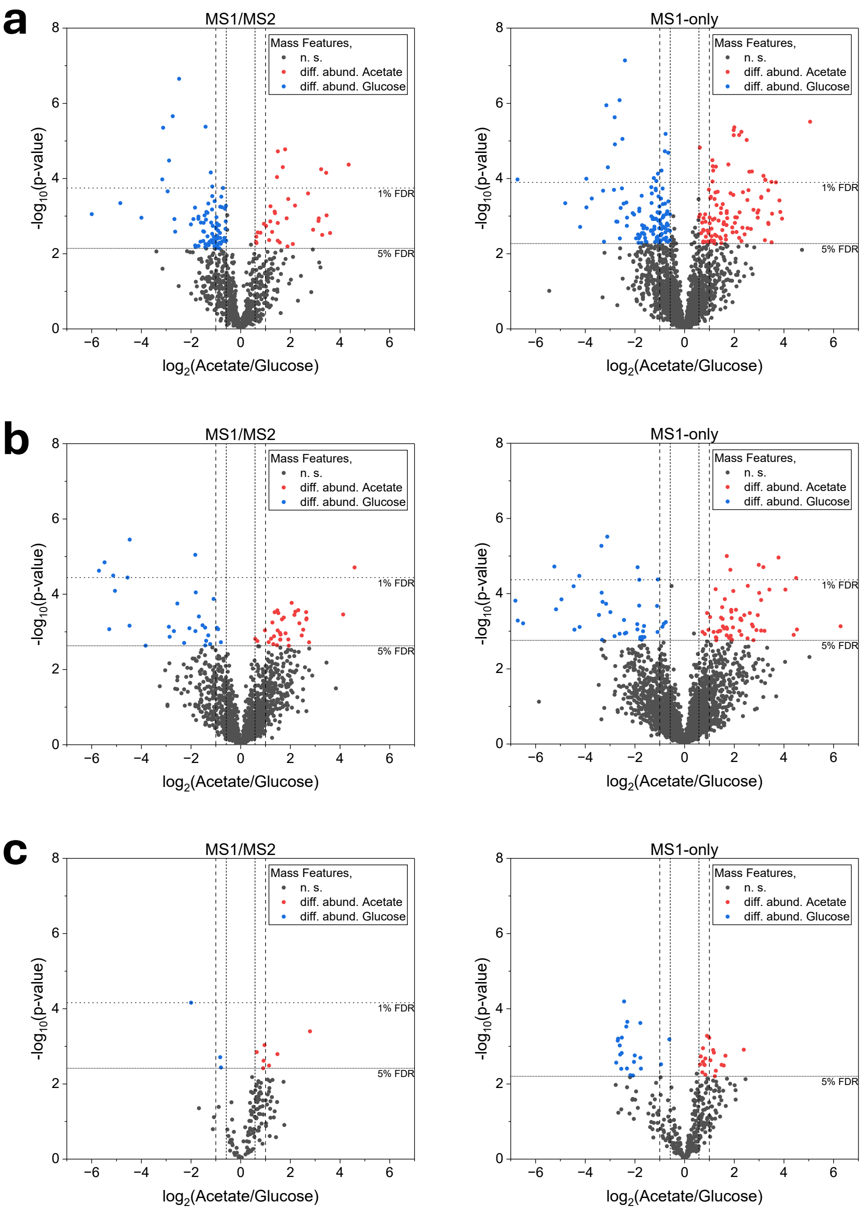


**Supplementary Figure 10**: **Superiority of the MS1-only approach using different experimental setups**. Shown are volcano plots at the mass feature level (no missing values). Left plots: MS1/MS2 approach; right plots: MS1-only approach. **a** 60-min gradient (C4 column) utilizing FAIMS (samples enriched in proteoforms suitable for TDP by solid-phase extraction). **b** 90-min gradient (MabPac column; samples enriched in proteoforms suitable for TDP by solid-phase extraction) without utilizing FAIMS. **c** 30-min gradient (full lysate without small proteoform enrichment, reduced and alkylated) without FAIMS using an MabPac column. Colored dots: significantly differentially abundant mass features more abundant in glucose (blue) or acetate (red) conditions. Horizontal lines: 5% and 1% FDR cutoffs (Benjamini-Hochberg correction); vertical lines: $\log_2$ fold changes of ±0.58 and ±1.

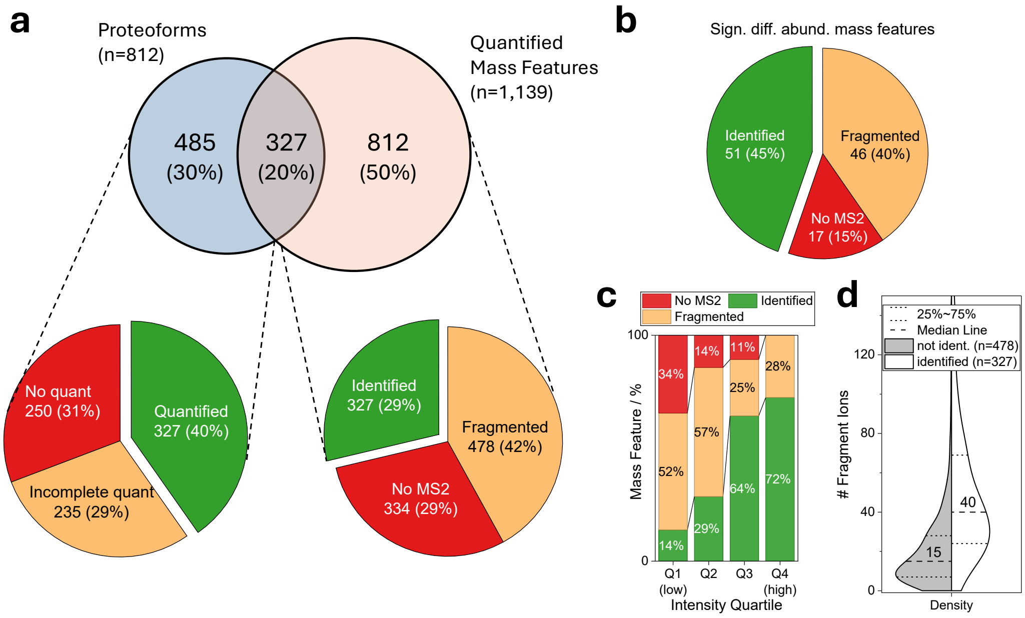


**Supplementary Figure 11**: **Factors limiting the overlap between quantified mass features and proteoform identifications in an MS1-only/targeted MS2 approach utilizing standard MS2 settings**. Significantly differentially abundant proteoforms were targeted. The same MS2 settings as for a classical DDA MS1/MS2 method were utilized. **a** Overlap between quantified mass features and identified proteoforms (Venn diagram) and factors why identified proteoforms could not be quantified (left pie chart), or quantified mass features could not be assigned to proteoform identifications (right pie chart). **b** Outcome of targeted MS2 acquisition of significantly differentially abundant mass features, showing the fractions that were identified, fragmented but not identified, or not selected for fragmentation. **c** Proteoform identification rate of data-dependent targeted significantly differentially abundant mass features as a function of mass feature intensity. **d** Distribution (scale: count) of fragment ion numbers in MS2 spectra that either resulted in successful proteoform identifications (labeled as identified) or remained unidentified (not ident.). The numbers represent the median values.

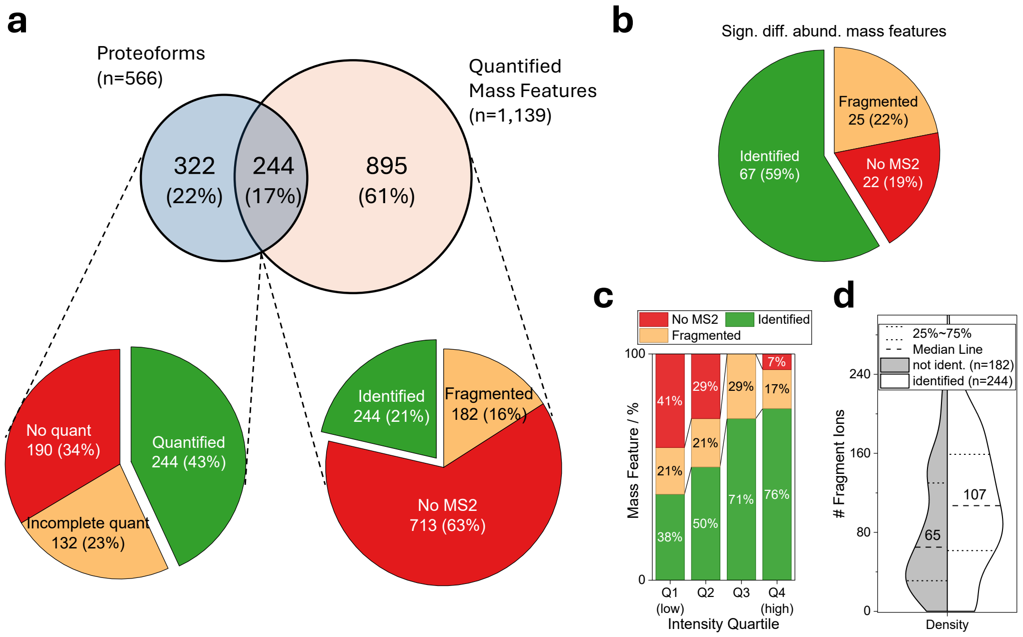


**Supplementary Figure 12**: **Factors limiting the overlap between quantified mass features and proteoform identifications in an MS1-only/targeted MS2 approach utilizing high-quality MS2 settings**. Significantly differentially abundant proteoforms were targeted. Compared to standard acquisition, MS2 spectra were acquired at higher resolution (120k) with longer maximum injection times (246 ms) to enhance fragment ion detection. In addition, two fragmentation spectra were acquired per precursor using HCD (30% NCE) and CID (35% NCE), respectively. **a** Overlap between quantified mass features and identified proteoforms (Venn diagram) and reasons why identified proteoforms could not be quantified (left pie chart) or quantified mass features could not be assigned to proteoform identifications (right pie chart). **b** Outcome of targeted MS2 acquisition of significantly differentially abundant mass features, showing the fractions that were identified, fragmented but not identified, or not selected for fragmentation. **c** Proteoform identification rate of data-dependent targeted significantly differentially abundant mass features as a function of mass feature intensity. **d** Distribution (scale: count) of fragment ion numbers in MS2 spectra that either resulted in successful proteoform identifications (labeled as identified) or remained unidentified (not ident.). The numbers represent the median values.

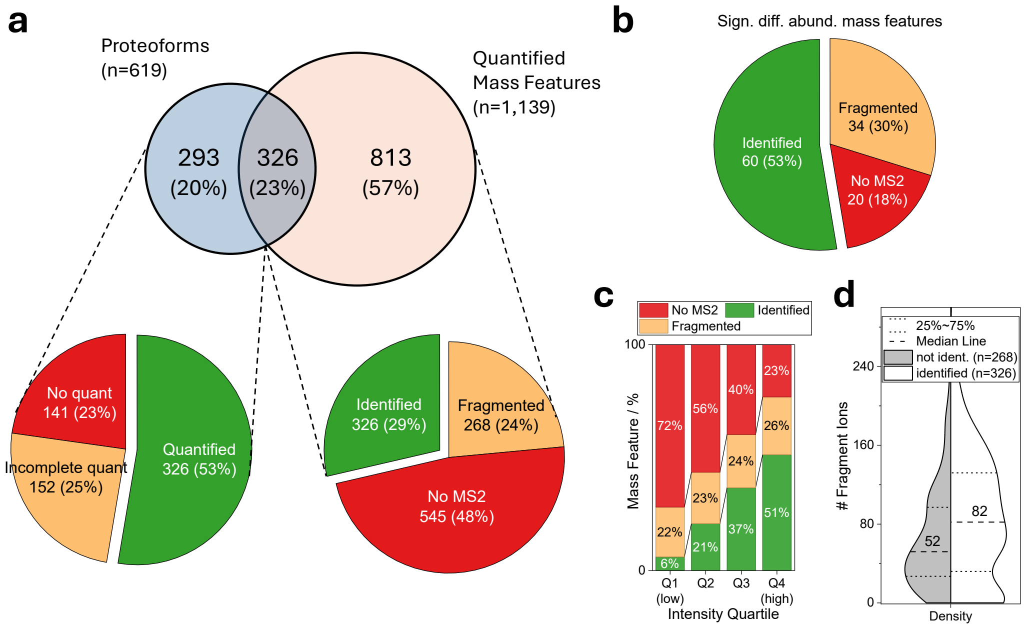


**Supplementary Figure 13**: **Factors limiting the overlap between quantified mass features and proteoform identifications in an MS1-only/targeted MS2 approach (targeting all quantified mass features) utilizing high-quality MS2 settings**. All quantified mass features were targeted. Compared to standard acquisition, MS2 spectra were acquired at higher resolution (120k) with longer maximum injection times (246 ms) to enhance fragment ion detection. In addition, two fragmentation spectra were acquired per precursor using HCD (30% NCE) and CID (35% NCE), respectively. **a** Overlap between quantified mass features and identified proteoforms (Venn diagram) and reasons why identified proteoforms could not be quantified (left pie chart) or quantified mass features could not be assigned to proteoform identifications (right pie chart). **b** Outcome of targeted MS2 acquisition of significantly differentially abundant mass features, showing the fractions that were identified, fragmented but not identified, or not selected for fragmentation. **c** Proteoform identification rate of data-dependent targeted quantified mass features as a function of mass feature intensity. **d** Distribution (scale: count) of fragment ion numbers in MS2 spectra that either resulted in successful proteoform identifications (labeled as identified) or remained unidentified (not ident.). The numbers represent the median values.

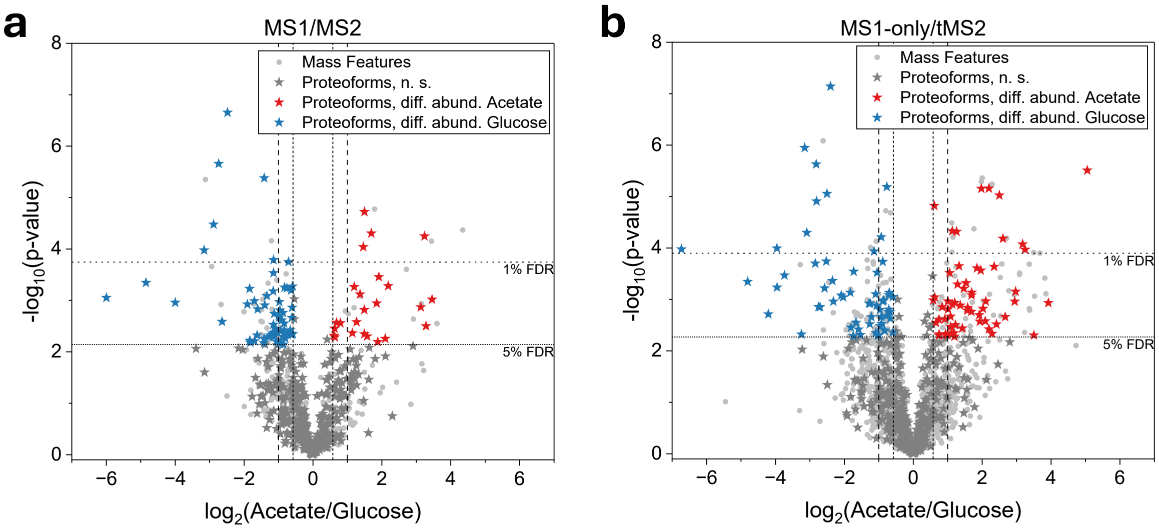


**Supplementary Figure 14**: **Quantified proteoforms obtained with the conventional MS1/MS2 workflow and the developed MS1-only/targeted MS2 workflow in a 60-min gradient**. **a** Conventional MS1/MS2 workflow. **b** MS1-only/targeted MS2 workflow. Statistical significance at the mass feature level was assessed using Welch's t-test with Benjamini-Hochberg correction. The stars represent proteoforms, and the dots represent mass features that do not result in proteoform identifications. Colored stars indicate significantly (FDR <5%) differentially abundant proteoforms with a fold change >1.5 (red, higher abundance in acetate; blue, higher abundance in glucose). Horizontal lines: 5% and 1% FDR cutoffs; vertical lines: $\log_2$ fold changes of ±0.58 and ±1.

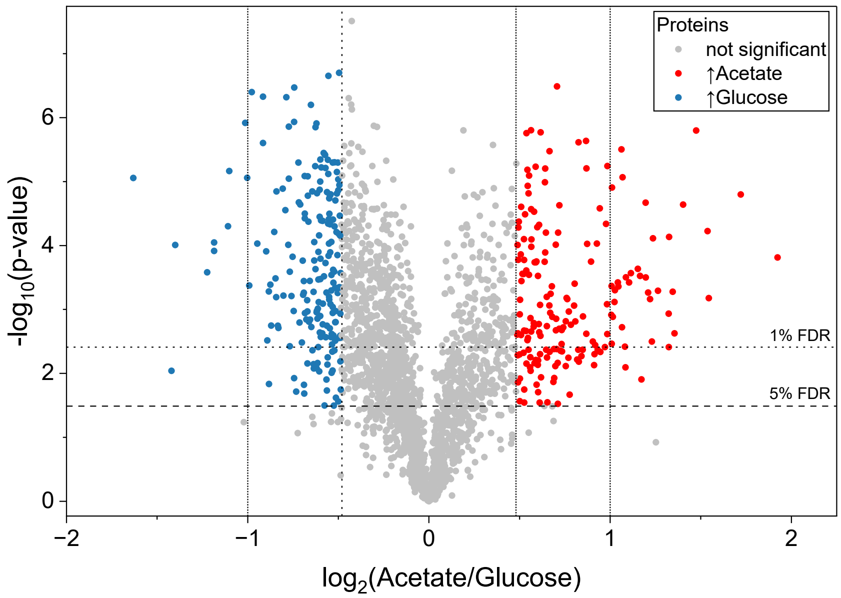


**Supplementary Figure 15**: Volcano plot of the reanalyzed BUP data. Each dot represents a protein, with colored ones are significantly differentially abundant (Welch's t-test with Benjamini–Hochberg correction). Horizontal lines: 5% and 1% FDR cutoffs; vertical lines: $\log_2$ fold changes of ±0.48 and ±1.

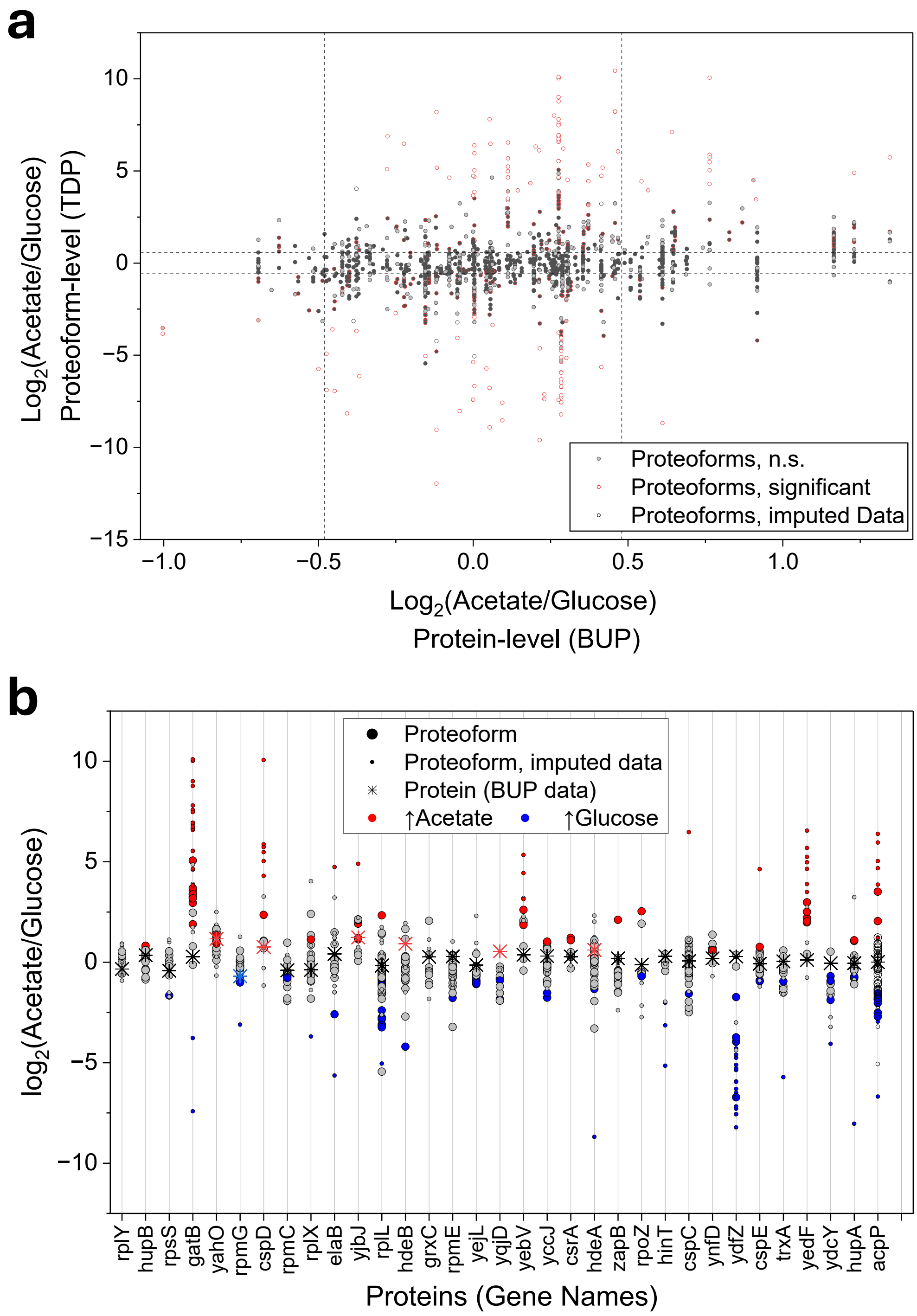


**Supplementary Figure 16:** Comparison of BUP protein-level and TDP proteoform-level quantification. **a** Proteins quantified in the BUP dataset and represented by at least two quantified proteoforms in the TDP dataset. Each dot represents a proteoform quantified by TDP. $Log_2$ fold changes from BUP protein-level quantification (x-axis) are plotted against those from TDP proteoform-level quantification (y-axis). **b** Comparison of protein- and proteoform-level quantification for proteins represented by at least 10 quantified proteoforms in the TDP dataset.

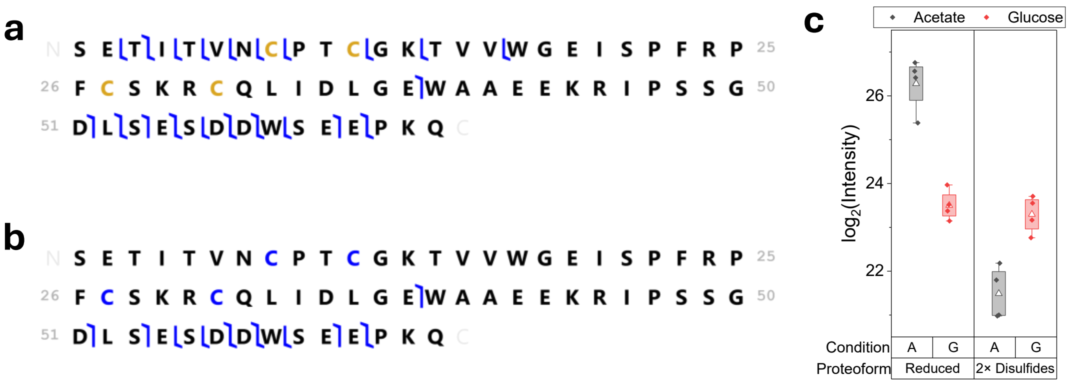


**Supplementary Figure 17:** Proteoforms of DNA gyrase inhibitor YacG (B1XC77) with opposite abundance between *E. coli* cultivated on glucose or acetate. **a** Proteoform with reduced cysteine residues, showing higher abundance under acetate conditions ($\log_2$ fold change: +2.78; mass error: 0.01 ppm; C-score: 586; p-score: 4.3e-53; residue cleavage: 33%; fragment explained: 55%). **b** Proteoform containing two disulfide bridges, showing higher abundance under glucose conditions ($\log_2$ fold change: −1.8; mass error: −2.53 ppm; C-score: 244; p-score: 4.9e-26; residue cleavage: 16%; fragment explained: 52%). **c** Normalized $\log_2$ intensities of the two proteoforms under acetate (A) and glucose (G) growth conditions. Boxplot: 25-75%, range within 1.5 IQR, triangle: mean.

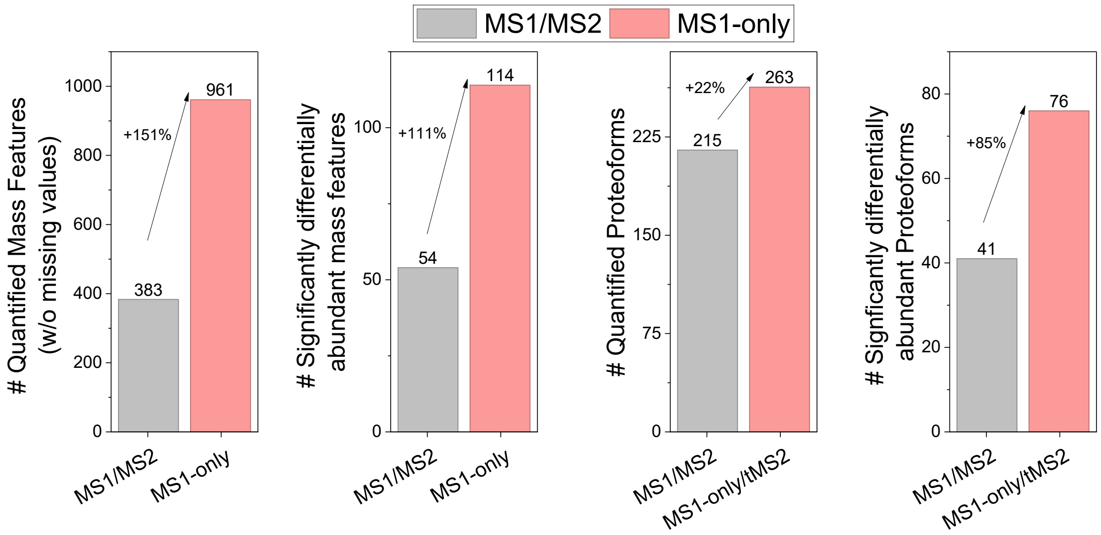


**Supplementary Figure 18**: **Filtering potential isotopic errors clearly preserved the trends observed in the original analysis**. After feature alignment, mass features with a mass difference of 1.00±0.02 Da and an apex retention time difference of less than 10 s were grouped, and only the most abundant feature in each group was retained. All other features were excluded. For matching quantified mass features to proteoform identifications, potential isotopic errors were also considered. Dataset: 30 min acquisitions. For tMS2, differentially abundant mass features were targeted, and two high-quality fragmentation spectra were acquired; note that no additional targeted experiments were performed, but the targeted results from the original analysis (see Figure 4) were used. The MS1-only/tMS2 workflow outperformed conventional MS1/MS2 acquisition, with the same trends observed in the original analysis (see Figure 3 and Figure 4).

### Supplementary Tables

**Supplementary Table 1**: Quantified mass features and assigned proteoforms of *Escherichia coli* cells cultivated with glucose or acetate. Statistical analysis was based on MS1-only acquisitions using a 60-min gradient and multiple targeted and untargeted MS2 acquisitions. Mass features that were detected in all four biological replicates of at least one condition were used for quantification, and data imputation for missing values was applied. Excel file.

**Supplementary Table 2**: Quantified proteins of a reanalyzed TMT-based BUP dataset of *Escherichia coli* cell cultivated with glucose or acetate and comparison with TDP proteoform-level results. Excel file.
